# A retrograde transit filter mediated by optineurin controls mitostasis in distal axons

**DOI:** 10.1101/2024.07.28.604753

**Authors:** Natalia A. Marahori, Barbara Gailer, Martina Schifferer, Tatjana Kleele, Anna Iatroudi, Shabab B. Hannan, Petros Avramopoulos, Stefan Engelhardt, Melike Lakadamyali, Monika S. Brill, Thomas Misgeld

**Affiliations:** Institute of Neuronal Cell Biology, Technische Universität München, Biedersteiner Straße 29, 80802 Munich, Germany; Institute of Advanced Study (IAS), Technische Universität München, Lichtenbergstraße 2 a, 85748 Garching, Germany; German Center for Neurodegenerative Diseases (DZNE), Feodor-Lynen-Straße 17, 81377 Munich, Germany; Munich Cluster of Systems Neurology (SyNergy), Feodor-Lynen-Straße 17, 81377 Munich, Germany; Institute of Pharmacology and Toxicology, Technische Universität München, 80802 Munich, Germany; German Center for Cardiovascular Research (DZHK), partner site Munich Heart Alliance, 80802 Munich, Germany; Department of Physiology, Perelman School of Medicine, University of Pennsylvania, Philadelphia, PA, 19104, USA

**Author notes:** Institute of Biochemistry, Eidgenössische Technische Hochschule (ETH) Zürich, 8093 Zürich, Switzerland. Jan and Dan Duncan Neurological Research Institute, Baylor College of Medicine, 1250 Moursund St., Houston, TX 77030.

**Keywords:** mitochondria, neuromuscular junction, axonal transport, autophagy, mitophagy, ALS, synapse

## Abstract

The mechanisms of mitochondrial homeostasis in neurons—‘mitostasis’—are enigmatic and disease-prone, given a neuron’s large synaptic pool of mitochondria. To better understand mitostasis in mature mouse motor axons, we used ex vivo optical pulse-chase measurements of mitochondrial volume flux and identified a reiterative degradation system near presynaptic terminals and distal paranodes, which accounts for 75% of mitochondrial turnover in synapses. This distal filter system captures dysfunctional mitochondria from the retrograde stream and redirects them for lysosomal degradation. It depends on optineurin, a motor neuron disease-related mitophagy adaptor, but not on PINK1 and parkin, implicating a non-canonical mitophagy pathway. In presymptomatic motor neuron disease models, where retrograde transport is disrupted, the fraction of removed mitochondria is increased. Thus, we identify a new mitostasis system in distal axons with a cascade of checkpoints for local mitophagy, which normally maintains mitochondrial mass balance but is disrupted early in degenerative axonopathies.

## MAIN

Balancing the number of mitochondria and maintaining their quality (‘mitostasis’) is challenging for neurons with their extended morphology (Misgeld & Schwarz, 2017). This is especially true for long and branched projection neurons such as those affected by Parkinson’s disease (PD) or amyotrophic lateral sclerosis (ALS). Two processes—mitochondrial transport and degradation—are key to maintaining neuronal mitostasis, as evidenced by mutations in the molecular pathways mediating these processes in familial forms of PD and ALS (Abeliovich & Gitler, 2016; Evans & Holzbaur, 2019a; De Vos et al., 2017; Killackey et al., 2020). Yet, the interplay of these two processes and their specific contribution to mitostasis remain to be elucidated—especially because previous attempts to quantitatively account for mitochondrial turnover in axons have failed, hinting at hitherto hidden sites of mitochondrial degradation (Misgeld & Schwarz, 2017). Indeed, the cellular location of key mitochondrial degradation pathways in mammalian neurons *in vivo* remains a matter of debate, even though it has become clear that, *in vitro,* these pathways consist of highly compartmentalized steps that can be spaced along neurites and are linked together by transport (Evans & Holzbaur 2019b; Maday, 2016; Hill & Colon-Ramos, 2020). In particular, we lack a comprehensive view of how mitochondrial health is maintained at synapses, the major functional sites of a neuron that are also the most prone to degeneration due to their high metabolic activity and distal localization (Pekkurnaz & Wang, 2022; Palikaras & Tavernarakis, 2020).

Mitochondrial degradation occurs on the scale of individual proteins, small sections of mitochondria, and entire organelles (Ashrafi & Schwarz, 2013). Mitophagy is an organelle-selective form of autophagy and follows a distinct molecular pathway compared to unselective ‘bulk’ autophagy: during mitophagy, defective mitochondria are labeled by ubiquitin via proteins such as PINK1/parkin—gene products related to PD—and then ‘swallowed’ by an autophagosome for subsequent lysosomal degradation (Stavoe & Holzbaur, 2019). Linking ubiquitin chains and the autophagosomal membrane requires mitophagy receptors, some of which are defective in ALS, like optineurin and SQSTM1 (Cirulli et al., 2015; Maruyama et al., 2010; Palikaras et al., 2018). Notably, several potentially redundant mitophagy pathways exist (Evans & Holzbaur 2019b). The *in vivo* roles of these pathways are still unresolved, and their existence in axons or synapses remains disputed. For instance, it is unclear whether neurons use the same canonical quality control mechanisms for peripheral mitochondria as for somatic mitochondria (Evans & Holzbaur 2019b). Some models posit a peripheral engulfment of mitochondria in neurites by ‘bulk’ autophagy, which is followed by the retrograde transport of these degradative organelles to the soma for maturation and fusion with a lysosome (Maday, 2016; Neisch et al., 2017), while other models imply that selective mitophagy may occur locally in the periphery (Ashrafi et al., 2014). Other lines of argument dispute the existence of distal mitochondrial degradation (Cai et al., 2012; McWilliams et al., 2018) by mature axonal lysosomes (Han et al., 2020; Lie et al., 2021) and suggest that mitophagy is ultimately somatic and hence a chief ‘raison d’etre’ for retrograde shuttling of mitochondria (Sheng & Cai, 2012; Sheng, 2014). Given the implied compartmentalization of autophagy-related processes, it seems conceivable that different neuronal compartments use different pathways for mitochondrial degradation, a question that is significant as, depending on the answer, different patterns of axon degradation would be expected, and distinct interventional strategies could be pursued to modulate neuronal mitophagy. Many aspects of the ongoing debates likely relate to the specific cell culture systems used. So, to settle the question of whether mitophagy occurs at distal sites in fully differentiated neurons, as well as to elucidate the cell biological and molecular mechanisms of distal axonal mitostasis, we choose the projections of mouse alpha motor neurons as model axons due to their experimental accessibility (Kerschensteiner et al., 2008) and relevance in mitophagy-related diseases such as ALS. We identified distal perinodal regions, especially the synaptic ‘exit point’ of the NMJ, as local ‘checkpoints’ for the quality control of retrogradely moving mitochondria, where defective organelles are removed by optineurin-dependent but PINK1/parkin-unrelated mitophagy.

## RESULTS

### Most mitochondria delivered to distal motor axon branches vanish in synapses

Axonal transport measurements from adult animals show that *in vivo*, neurons appear to deliver more mitochondria anterogradely toward the axonal periphery than return retrogradely to the soma (Misgeld et al., 2007; Misgeld and Schwarz, 2017; Xu et al., 2017; Chen et al., 2016; Sorbara et al., 2014). Despite the surplus, mice typically maintain a stable mitochondrial volume in mature neuromuscular junctions (NMJs) that show limited growth (≥ 6 weeks; Marinkovic et al., 2012; Lichtman et al., 1987) and do not seem to transfer mitochondria in discernible numbers from axons to sheathing glia (Misgeld et al., 2007; Misgeld and Schwarz, 2017). This suggests that a large portion of peripheral mitochondria is degraded along the axon or in synaptic terminals, implying the existence of an unknown site of mitochondrial degradation on a massive scale. Alternatively, however, anterogradely moving mitochondria may simply be smaller than retrogradely moving ones, resulting in a balanced net mitochondrial ‘volume flux’.

We set out to disambiguate these possibilities by establishing optical ‘pulse-chase’ experiments in motor neurons of acute ex vivo nerve-muscle explants (**Fig. 1a**) from adult *Thy1*-mito-XFP mice (i.e. transgenic mice, where various mitochondrially-targeted fluorescent proteins are expressed under a neuronal promoter; see **Extended Data Fig. 1a–d** for new lines generated for this study; **Methods** for established lines; Misgeld et al., 2007; Marinkovic et al., 2012; Breckwoldt et al., 2014). These allow for linking transport flux measurements to single mitochondria volume estimates (Misgeld et al., 2007; Marinkovic et al., 2012). First, we corroborated the validity of our widefield imaging for volume estimates near the optical diffraction limit (which mitochondrial diameter approaches) using correlated Airyscan microscopy, which can resolve sub-mitochondrial morphology (i.a. Kondadi et al., 2020; **Extended Data Fig. 2**).

**Fig. 1:**
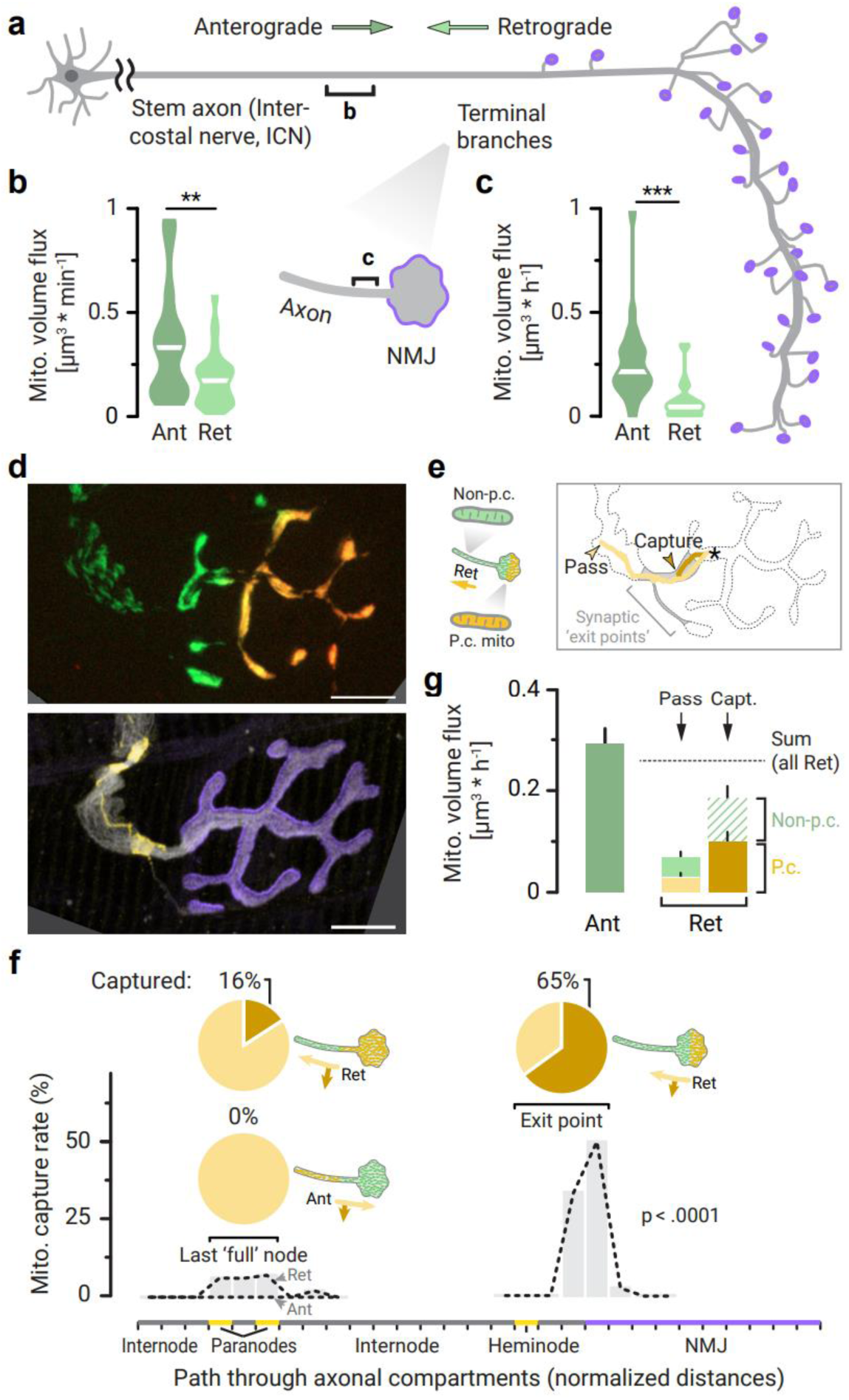
Retrogradely moving mitochondria are detained in distal motor axon branches and vanish. **a**, Schematic illustrating the geometry of mature α-motor neuron projections. Brackets show live imaging positions in acute intercostal nerve-muscle explants, where measurements **b** and **c** were taken (NMJ: neuromuscular junction). **b, c** Anterograde (‘Ant’) and retrograde (‘Ret’) mitochondrial volume flux rates in **b**, stem axons (29 axons from 14 *Thy1*-mito-XFP transgenic mice; ‘XFP’ were various fluorescent tags, see Methods) and in **c**, distal axon branches with synaptic terminals (25 axons, 16 *Thy1*-mito-XFP mice). **d**, Top: Widefield projection of presynaptic motor terminal showing Dendra-tagged mitochondria that were photoconverted (p.c.) from green to red fluorescence at the NMJ tip (*Thy1*-mito-Dendra line). Bottom: Post-hoc staining of the same axon terminal with marked nodes of Ranvier (paranodal components, anti-Caspr-antibody, yellow), acetylcholine receptors (α-bungarotoxin, magenta), neuronal microtubules (βIII-tubulin-antibody, gray). **e**, Left: Schematic illustrating the experimental setup (p.c. synaptic mitochondria that initiated retrograde movement were tracked on the non-p.c. background). Right: Trackings of two p.c. mitochondria. Star: starting point of tracks; dark orange: permanent stop (‘captured’ mitochondrion) in synaptic exit point (gray area), light orange: continuation into axon (‘pass’). **f**, Graph indicates probability that p.c. mitochondria stop permanently while traversing through internodes, paranodes/heminode, and NMJ (some compartments were sectioned along their length into bins; see Methods). Various mitochondrial populations were tracked in different locations (p.c. sites and directions of mitochondrial transport indicated in schematics). Pie charts: proportion of tracks that ended in ‘capture’ (%, dark orange) at either the last ‘full’ node of Ranvier or the synaptic exit point (n ≥ 19 mitochondria from ≥ 4 axons, ≥ 3 *Thy1*-mito-Dendra mice per group for ‘full’ nodes; n = 40 mitochondria for exit points, analyzed from the movies in **g**). **g**, Mean mitochondrial volume flux rates at synaptic exit points. The sum of ‘captured’ and ‘passing’ flux rates is indicated by a dotted line (see **Extended Data Fig. 3c** for individual data spread). Color code indicates p.c. (orange) and non-p.c. (green) populations, bar graphs were stacked to visualize the sum of both groups; hatched: non-p.c. ‘captured’ flux was estimated as described in Methods (n = 32 NMJs, 20 *Thy1*-mito-Dendra mice). Scale bars, 10 μm. Violin plots in **b,c** show data distribution and median; bars in **f** shown means per bin; bar graph in **g** mean ± s.e.m. Mann-Whitney test was used to determine significance in **b**,**c**: ∗∗∗p < 0.001; ∗∗p < 0.01; a χ²-test in **f** (‘Ret’ synaptic exit point p < 0.001; ‘Ant’ and ‘Ret’ last node of Ranvier p ≥ 0.05). See also **Supplementary Movies 1–4**

We then determined the volume flux of neuronal mitochondria in two locations along motor axons using time-lapse recordings: in the proximal stem of the axon in the intercostal nerve (ICN) before peripheral branches diverge off (Lu et al., 2009) and in the most distal terminal branches, beyond all branch points, just before the axon enters the NMJ (**Fig. 1a**–**c**). We indeed found that significantly more mitochondrial volume was transported anterogradely than retrogradely in ICN axons. This discrepancy was even bigger for mitochondrial volumes than for numbers, as retrogradely moving mitochondria were—if anything—smaller than anterograde moving ones (**Extended Data Fig. 2e,i**). Notably, this discrepancy was even more dramatic in terminal branches near the NMJ (**Fig. 1b,c**; ICN 1.9-fold vs. NMJ 3.4-fold difference from averaged transport rates, 2.2 and 29 total ‘axon imaging hours’, respectively).

### Most synaptic mitochondria that initiate retrograde movement are captured before exiting the NMJ

Since we observed an apparent overflow of mitochondria even at the most distal synaptic sites, we hypothesized that mitochondria must be removed within the confined compartment of the NMJ. To disambiguate whether removal involved the stationary pool or rather a hitherto unidentified dynamic class of mitochondria that traffics within the synapse, we devised an assay to partially label the synaptic pool of mitochondria using *Thy1*-mito-Dendra mice (**Extended Data Fig. 1e–h**; cf. Marinkovic et al., 2012; Magrane et al., 2014). We photoconverted mitochondria within the distal half of NMJs from green to red fluorescence (**Fig. 1d**). This pulse-chase approach (Marinkovic et al. 2012; Pham et al., 2012) allowed us to follow single mitochondria by time-lapse imaging against a dense stationary population of organelles, as they dispatched from the photoconverted pool and initiated retrograde movement. We then tracked these retrogradely moving mitochondria and mapped the track coordinates to anatomical landmarks after fixation (for used labels, see **Methods**; **Fig. 1e,f** and **Supplementary Movie 1**). Strikingly, we observed two distinct behaviors: while some mitochondria left the synaptic compartment and entered the axon (‘passed’ mitochondria, 35%), others stalled (‘captured’, 65%) and stayed in the NMJ for the rest of the observation time until the mitochondrion either disappeared or the movie ended (**Fig. 1f**; n = 40 mitochondria observed from 32 ‘NMJ-imaging hours’; **Extended Data Fig. 3a**). The probability of capture was highest in the transition area separating the NMJ and the distal-most internode, close to the terminal heminode (termed synaptic ‘exit point’ hereafter, **Fig. 1f**; χ²-test, p < 0.001). The overall volume flux of captured mitochondria (once corrected for the partial photoconversion of the NMJs; **Extended Data Fig. 3b**) accounted for the difference between the anterograde and retrograde volume flux, suggesting that captured mitochondria constituted the aforementioned ‘missing’ population (**Fig. 1g**; Ant. 0.28 ± 0.03 µm³/h, Ret. passed 0.07 ± 0.01 µm³/h, captured 0.19 ± 0.04 µm³/h; mean ± s.e.m.; n = 32 NMJs from 20 mice; also replotted in **Extended Data Fig. 3c**).

### Retrogradely moving mitochondria are selectively captured at synaptic exit points and distal nodes

Since the exit point is close to the last heminode, a specific nodal site, we wondered if any node of Ranvier could ‘capture’ retrogradely moving mitochondria. For this, we tracked photo-converted mitochondria as they moved across preterminal nodes (i.e., ‘full’ nodes, which are located upstream of the heminodal exit point; **Fig. 1f**), as well as in the stem axons. Occasionally, we observed capture of retrogradely moving mitochondria at the most distal ‘full’ node, albeit with a lower frequency than at the terminal heminode (**Fig. 1f**, **Supplementary Movie 2**; 16% probability of capture in distal ‘full’ nodes vs. 65% in heminode; p ≥ 0.05, χ^2^-test; n = 69 mitochondria observed from 9 total ‘axon imaging hours’ vs. NMJ measurements detailed above). In contrast, we did not encounter the capture phenomenon at stem axon nodes (**Supplementary Movie 3**, 0% capture, n ≥ 30 mitochondria observed from 3 total ‘imaging hours’, 3 axons from 3 *Thy1*-mito-Dendra mice). Moreover, anterogradely moving mitochondria were not captured at distal nodes (**Fig. 1f**, **Supplementary Movie 4**; 0% capture, n = 19 mitochondria observed from 6 total ‘axon imaging hours’), excluding unspecific causes for captures in the retrogradely moving population, such as the (bidirectional) reduction of motility around nodes of Ranvier (Ohno et al., 2011; Misgeld et al., 2007) or (symmetric) spatial constraints due to axon caliber constriction around nodes (**Extended Data Fig. 3d**; axon diameter, 54 ± 6% of preceding internode in anterograde vs. 43 ± 3% in retrograde direction; p ≥ 0.05, Mann-Whitney-U test; also **Extended Data Fig. 3e**), instead pointing to a selective mechanism that only acts on specific mitochondria.

Mitochondria that were later captured slowed down when approaching the exit point while initially moving at the same speed during their transport ‘runs’ compared to ‘passing’ mitochondria (**Extended Data Fig. 3f**; significant reduction of average run speed of captured mitochondria in exit point by 56%, compared to passed mitochondria; n = 49 mitochondria). In addition, ‘captured’ mitochondria were of more spherical shape than passing mitochondria (**Extended Data Fig. 3g**), potentially suggesting a dysfunctional state (Nikic et al., 2011; Zick et al., 2009; Ferree et al., 2013) or emergence from a distinct fission event (Kleele et al., 2021). To determine whether capture relates to mitochondrial functionality, we used a photo-caged mitochondrial depolarizing agent (mito-photo-DNP; Chalmers et al., 2012). This allowed us to reduce the membrane potential of mitochondria in individual NMJs via photoactivation (**Fig. 2a,b**). Photo-depolarization indeed amplified the proportion of captured mitochondria at the exit point, compared to a ‘sham’ experiment (**Fig. 2c,d**; capture fraction increased 2.5-fold, n ≥ 12 mitochondria per group from ≥ 6 ‘NMJ imaging hours’, p = 0.003, χ²-test).

**Fig. 2:**
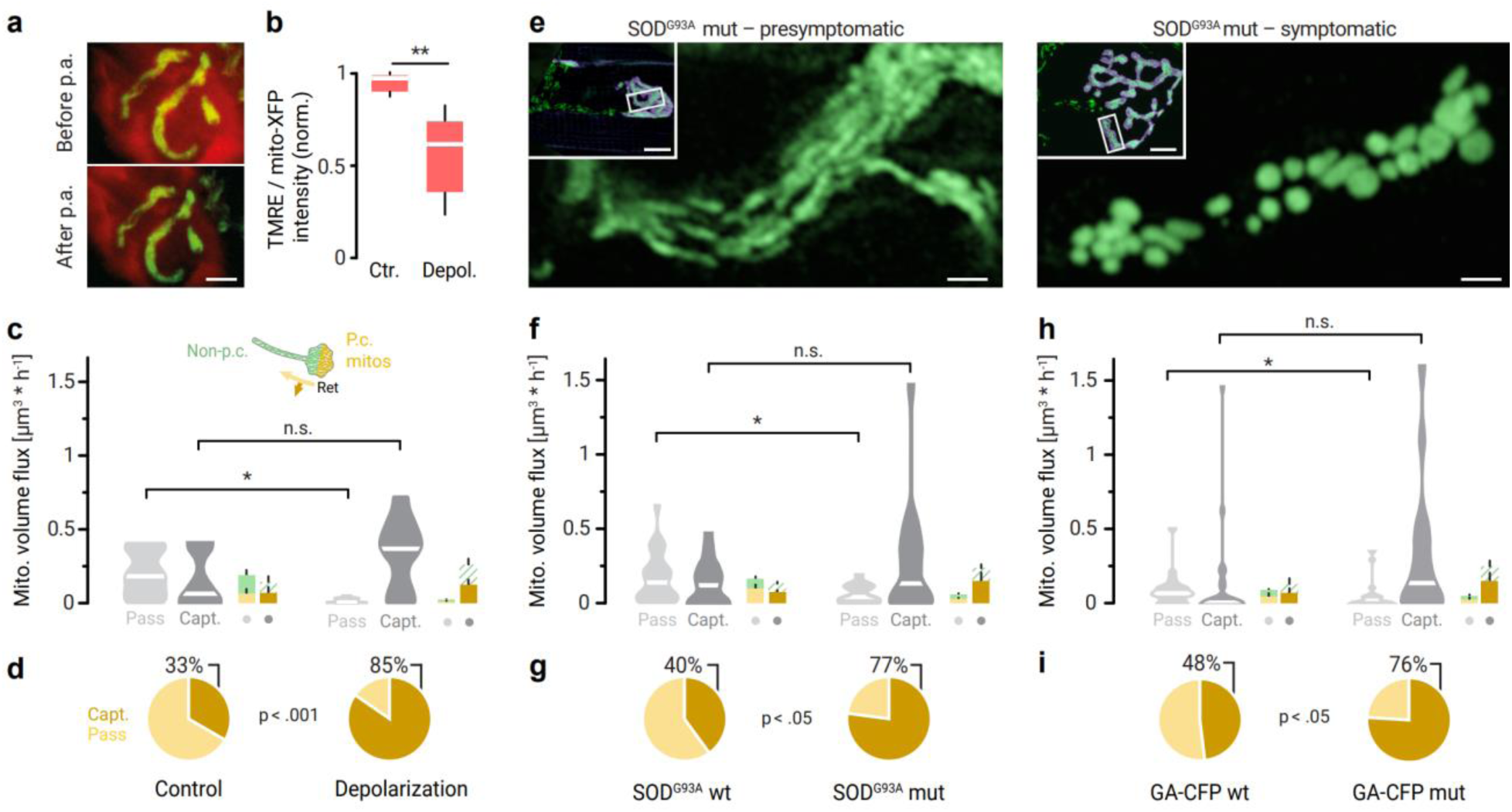
Mitochondrial damage increases the fraction of captured mitochondria. **a**,**b**, Depolarization of presynaptic mitochondria in acute nerve-muscle explants after incubation and photoactivation (p.a.) of photo-mito-DNP by using a local UV light pulse (ChAT-Cre^+^ × ROSA-mito-GFP mice, see Methods). **a**, Presynaptic mitochondria (in green) were labelled with the mitochondrial membrane potential dye TMRE (in red). Top: NMJ before p.a. of the mito-DNP compound, Bottom: NMJ after p.a. **b**, Fluorescence intensity of TMRE after p.a. of photo-mito-DNP (normalized to the start of the experiment, Ctr: only UV; n ≥ 5 NMJs from 1 mouse per group). **c**,**d**, Depolarized mitochondria are preferentially captured at synaptic exit points (Left: control group without depolarization of synaptic mitochondria; Right: experimental group with mitochondria depolarized using photo-mito-DNP activation; see Methods) **c**, Mitochondrial volume flux rates at synaptic exit points (n ≥ 6 NMJs, ≥ 4 *Thy1*-mito-Dendra mice per group). The box plots show mitochondrial volume flux rates of all passing (light grey) and captured (dark grey) mitochondria. Miniature bar graphs show stacked mean values for both p.c. (orange) and non-p.c. populations (green) measured in the passing (light shades) and captured (dark shades) populations (see Fig. 1 **d**–**g** for experimental setup). **d**, Pie charts represent the respective fractions of p.c. mitochondria that were ‘captured’ (%, dark orange) synaptic exit points (n ≥ 12 mitochondria per group, from the movies in **c**). **e**, Single optical sections of presynaptic mitochondria (green, Airyscan microscopy) in motor axon terminals of ALS-related mice (SOD^G93A^ ^+/-^ mutation); Left: Triangularis sterni muscle of a presymptomatic, 2-week-old mouse, Right: Fragmented mitochondria in a 14-week-old, symptomatic mouse (Insets: boxes indicate positions of the images inside maximum projections of the axon terminals; α-bungarotoxin, magenta; *Thy1*-mito-Dendra transgene, green). **f**,**g**, Synaptic mitochondria in presymptomatic SOD^G93A^ ^+/-^ mice are preferentially captured at synaptic exit points (Right: experimental group with two-week old SOD^G93A^ ^+/-^ (“mut”) × *Thy1*-mito-Dendra mice; Left: group of healthy SOD^G93A-/-^ (“wt”) × *Thy1*-mito-Dendra littermates; n ≥ 13 mice per genotype). **f**, Mitochondrial volume flux at synaptic exit points (Legend: as in **c**) (n ≥ 19 NMJs per genotype). **g**, Fractions of p.c. retrogradely moving mitochondria that were captured at synaptic exit points (n ≥ 48 mitochondria per genotype). **h**,**i**, Mitochondria in presymptomatic GA-CFP ^+/-^ mice are preferentially captured at synaptic exit points (Right: experimental group with 5-week-old GA-CFP ^+/-^ (“mut”) × *Thy1*-mito-Dendra mice; Left: group of healthy GA-CFP ^-/-^ (“wt”) × *Thy1*-mito-Dendra littermates; n ≥ 11 mice per genotype). **h**, Mitochondrial volume flux at synaptic exit points (Legend: as in **c**) (n ≥ 19 NMJs per genotype). **i**, Fractions of p.c. mitochondria being captured at synaptic exit points (n ≥ 26 mitochondria per genotype). Scale bars in **a** 10 μm; **e** 1 µm (10 µm in insets). Box plot in **b** shows median, quartile 1 - quartile 3, whiskers 95^th^ percentiles. Bar graphs in **c,f,h**: mean ± s.e.m.; Violin plots in **c,f,h** show data distribution with median. Mann-Whitney test was used to determine significance in **b**,**c**,**f**,**h**: ∗∗p < 0.01; ∗p < 0.05; n.s. non-significant; a χ²-test was used in pie charts (**d**, p = 0.003; **g**, p < 0.001; **i**, p = 0.047).

We next explored whether disease-related damage would also increase the captured fraction. We tested this in two mouse models of ALS (namely SOD^G93A^; Gurney et al., 1994; and GA-CFP; Schludi et al., 2017). We focused on the pre-symptomatic stages in SOD^G93A^ × *Thy1*-mito-Dendra mice at two weeks of age before the onset of transport deficits in the ICN and degeneration of the NMJ (Marinkovic et al., 2012; **Fig. 2e**). Indeed, in SOD^G93A^ expressing mice, the mitochondrial capture fraction in exit points was increased by 1.9-fold compared to healthy littermate controls (**Fig. 2g**; n ≥ 48 mitochondria from ≥ 19 ‘NMJ imaging hours’ per genotype, p < 0.001, χ²-test; furthermore, a reduction in ‘passing’ mitochondrial volume flux was observed, **Fig. 2f**). Similarly, in the dipeptide repeat protein-based C9orf72 mouse model (GA-CFP × *Thy1*-mito-Dendra), presymptomatic (5-6 weeks old) mice showed a ∼1.6-fold increased fraction of captured mitochondria (**Fig. 2i**; n ≥ 26 mitochondria from ∼19 NMJ imaging hours per genotype, p = 0.047, χ²-test; with reduction in ‘passing’ mitochondrial volume flux was observed, **Fig. 2h**).

In summary, we identified a specific mechanism where cascades of perinodal checkpoints (starting at the synaptic ‘exit point’ at the last heminode) filter out and remove damaged mitochondria from the retrograde motile pool as they transit towards the soma—a system that is employed in healthy mature axons and is adjusted in degenerative axonopathies related to ALS.

### Lysosomes are enriched in NMJ exit points, where mitochondria undergo lysosomal degradation

If mitochondria are captured at synaptic exit points, are they also locally degraded? To explore this, we intrathecally injected neonatal mice with an adeno-associated viral (AAV) vector expressing a ratiometric mitophagy indicator under a synapsin promotor (AAV9-*hSyn*-mito-Keima), thus visualizing mitochondria (neutral pH) in addition to lysosomes degrading mitochondrial material (‘mitolysosomes’, defined by an acidic environment that shifts Keima’s spectrum; Katayama et al., 2011). In adult motor axons, mitolysosomes were frequently encountered across the entire NMJ but were markedly enriched at synaptic exit points and near paranodes (**Fig. 3a**–**d**), mirroring the localization of mitochondrial capture sites. Similarly, organelles labeled with acidophilic vital dyes (which, i.a., label lysosomes; Song et al., 2008) accumulated at these locations (**Fig. 3e**–**h**). This labeling also coincided with electron-dense organelles, filled with a heterogeneous content characteristic of lysosomes in correlated volume electron microscopy (**Fig. 3i**–**l**). The majority (91% of 415 particles) of acidophilic dye-stained organelles were mitolysosomes (**Extended Data Fig. 4a–c**), which were not visible using regular mitochondrial markers (*Thy1*-mito-XFP; **Extended Data Fig. 4d,e**), indicating an actively digesting state of lysosomes or related endo-lysosomal organelles.

**Fig. 3:**
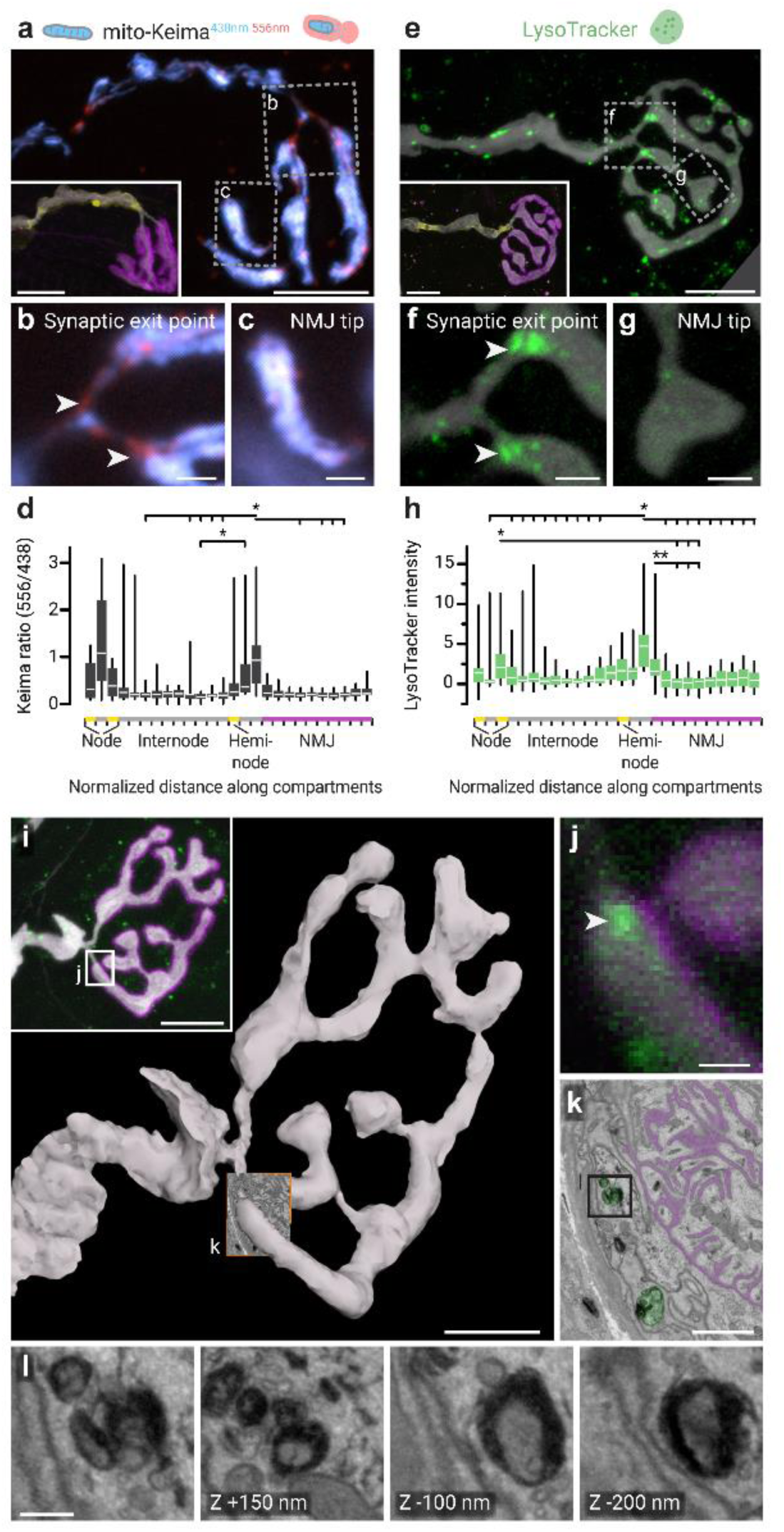
Accumulation of degradative organelles at mitochondrial ‘capture’ sites of motor axons. **a**–**d**, Mitolysosomes in AAV9-*hSyn*-mito-Keima expressing mice. **a**, Widefield projection of mito-Keima fluorescence after excitation with 438 ± 12 nm (shown in blue) and 556 ± 10 nm light (red). Inset: correlated staining (anti-Caspr, yellow; α-bungarotoxin, magenta; anti-βIII-tubulin, white). Boxes show positions of widefield images in **b**, Synaptic exit point with mitolysosome accumulations (high 556/438 ratio; arrowheads), and **c**, Tip of the axon terminal. **d**, Mito-Keima signal ratio at 556/438 nm excitation along different compartments in motor axon terminals (NMJ, magenta; heminode and paranodes, yellow; internode, gray; n = 14 axons, 7 *Thy1*-mito-CFP and C57BL/6 mice). **e**–**h**, Degradative organelles visualized by an acidophilic dye. **e**, Confocal projection of LysoTracker staining (green) in a YFP-expressing neuron (white, *Thy1*-YFP-16 line). Inset: correlated staining, anti-Caspr (yellow), α-bungarotoxin (magenta). Boxes show positions of single optical sections through the synaptic exit point (**f**) and tips (**g**), with LysoTracker accumulations (arrowheads) apparent within the exit point. **h**, LysoTracker fluorescence intensity inside axon terminals (normalized to internodes; n = 20 axons, 3 *Thy1*-YFP-16 mice). **i**–**m**, Correlative electron microscopy of LysoTracker-stained organelles. **i**, Three-dimensional rendering of a re-localized axon terminal, based on tracings in serial electron micrographs. Inset: LysoTracker staining (green) in a confocal projection of the corresponding axon terminal before fixation (α-bungarotoxin, magenta; *Thy1*-YFP-16, white). **j**, Confocal single optical section through the synaptic exit point showing a stained organelle accumulation (arrowhead); section position indicated by box in **i**. **k**, Ultrathin section through the targeted organelles (pseudo-colored green; post-synaptic folds: pseudo-colored magenta; section position shown in rendering, **i**). **l**, Micrograph of electron-dense organelles at the position of the box in **k** (micrograph on the very left; the other micrographs are taken at the same xy-position but from different sections in the 3D-stack; displacement in z-axis indicated in image corners). Scale bars in **a**,**e**,**I**,**k** 10 μm; **b**,**c**,**f**,**g** 2 µm; **j**,**l** 1 µm; **m** 200 nm. Box plots: median, quartile 1 - quartile 3, whiskers 95th percentiles. Statistical significance was determined using a Kruskal-Wallis-test (**d,h**) with Dunn’s multiple comparisons test (∗∗p < 0.01; ∗p < 0.05). See also **Extended Data Fig. 4**

To further define the relationship between stalling and autophagic digestions, we explored whether captured mitochondria were actually ‘bona fide’ mitochondria subsequently undergoing local mitophagy at exit points or if they were already enclosed in a pre-degradative organelle, such as an autophago-lysosome, during their transit to the exit point (Maday et al., 2012). We thus photoactivated the distal portion of NMJs in *Thy1*-mito-paGFP mice (injected with AAV9-*hSyn*-mitoKeima) and monitored the acidity of retrogradely moving photoactivated mitochondria at various locations along their route. Neither captured nor passing mitochondria colocalized with an acidic Keima-signal while moving, but captured mitochondria showed signs of acidication at the exit point (**Extended Data Fig. 4f–h**). Further, mitolysosomes had different shapes and kinetic parameters compared to motile mitochondria that were later captured (**Extended Data Fig. 4i–j**), and moved in a manner uncharacteristic of captured mitochondria, such as frequently attaching and detaching from exit points with bouts of fast bidirectional movement in-between (**Supplementary Movie 5**). These findings suggest that the captured organelles are bona fide, yet damaged mitochondria that are selectively recognized by an autophagic filter.

### Mitochondrial degradation, but not capture, is mediated by Atg7-related autophagy

We wondered whether ‘capture’ of mitochondria at synaptic exit points was directly related to the process of autophagosome formation. Specifically, gathering autophagosomes might ‘wait’ for damaged mitochondria at synaptic exit points and halt these organelles by locally initiating the engulfment process. Similar ‘autophagic filters’, where autophagosomes serve as gate-keepers for intracellular transport, have been suggested earlier at the axon hillock (Zaninello et al., 2020). To test whether such an autophagic filter exists at synaptic exit points, we blocked autophagy by deleting Atg7 selectively in cholinergic motor axons (ChAT-Cre^mut/wt^ × *Atg7*^fl/fl^, abbreviated below to *Atg7-*cKO; Komatsu et al., 2006; Komatsu et al., 2005). In *Atg7*-cKO mice, however, the volume flux rates of ‘passing’ mitochondria were not significantly increased when compared to littermate controls (**Fig. 4e**), refuting the hypothesis that an autophagic filter directly captures mitochondria at synaptic exit points.

**Fig. 4:**
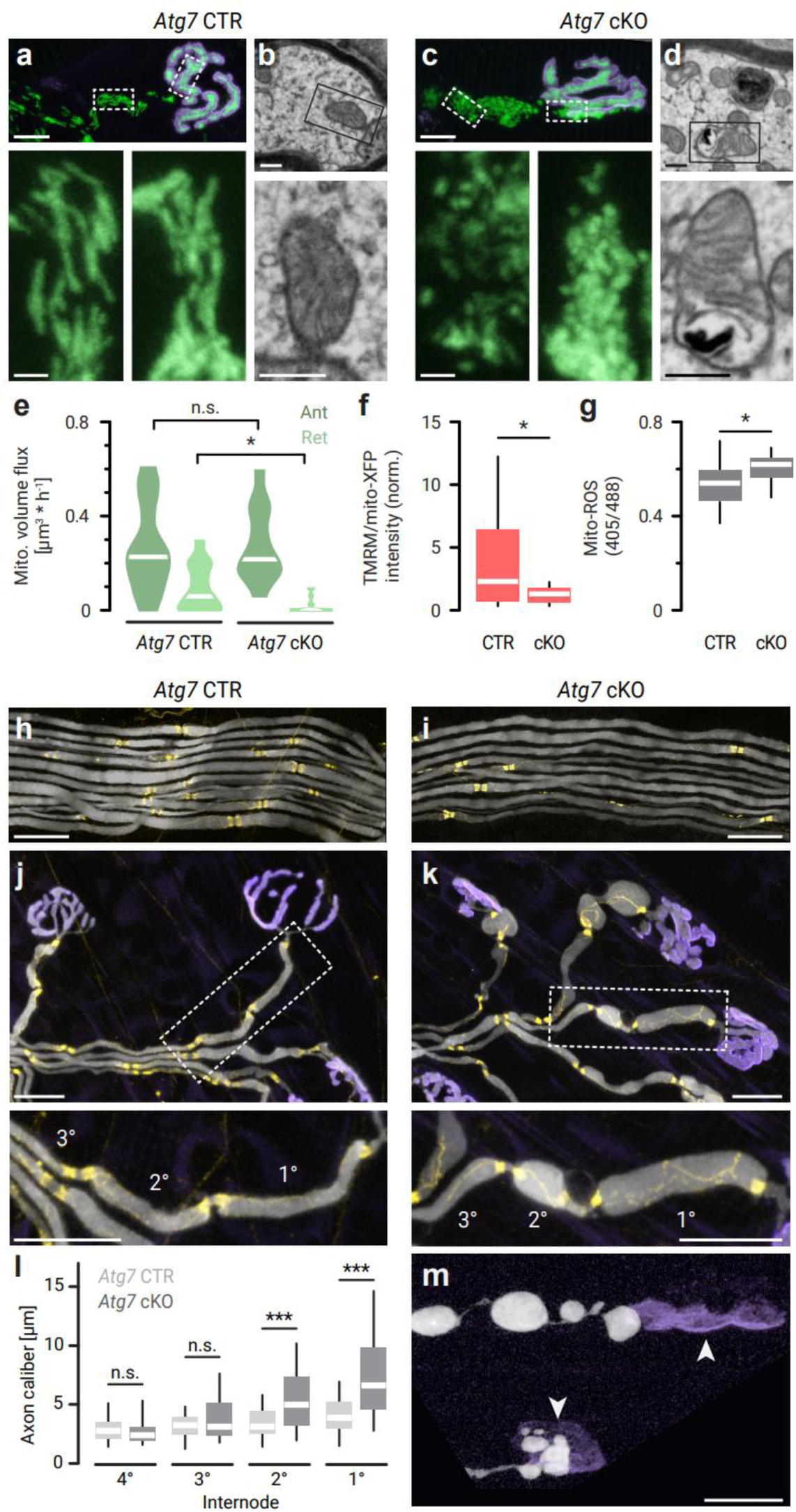
Damaged mitochondria accumulate in a distal-to-proximal manner in motor axons with impaired autophagy. **a**–**d**, Fragmented, spherical mitochondria with disrupted ultrastructure accumulate in neuromuscular junctions of 11-week-old Atg7 conditional-knockout mice (**a**,**b**: “Atg7 control (CTR)”, ChAT-Cre^wt/wt^ × *Atg7*^fl/fl^ × *Thy1*-mito-Dendra pooled with ChAT-Cre^mut/wt^ × *Atg7*^wt/wt^ × *Thy1*-mito-Dendra mice; **c**,**d**: “Atg7 cKO”, experimental group with ChAT-Cre^mut/wt^ × *Atg7*^fl/fl^ × *Thy1*-mito-Dendra littermates). **a**,**c**: Top row: confocal projections in acute nerve-muscle explants (green: *Thy1*-mito-Dendra, magenta: bungarotoxin), boxes indicate positions of single optical sections in the bottom row. **b**,**d**: Top: electron micrographs taken from axon terminals, boxes indicate positions of images in the bottom row. **e**, Mitochondrial volume flux rates in axon terminals of acute nerve-muscle explants measured in anterograde and retrograde direction (“Atg7 CTR”, ChAT-Cre^wt/wt^ × *Atg7*^fl/fl^ × *Thy1*-mito-Dendra pooled with ChAT-Cre^mut/wt^ × *Atg7*^wt/wt^ × *Thy1*-mito-Dendra mice; “Atg7 cKO”, experimental group with ChAT-Cre^mut/wt^ × *Atg7*^fl/fl^ × *Thy1*-mito-Dendra littermates; n ≥ 13 axons from ≥ 8 mice per group). **f**, Membrane depolarization of presynaptic mitochondria in acute nerve-muscle explants. Fluorescence pixel intensity of the mitochondrial membrane potential dye TMRM, normalized to a neuron-specific mitochondrial matrix label (mito-XFP; “CTR”, ChAT-Cre^wt/wt^ × *Atg7*^fl/fl^ × *Thy1*-mito-CFP × *Thy1*-YFP16 pooled with ChAT-Cre^mut/wt^ × *Atg7*^wt/wt^ × *Thy1*-mito-CFP × *Thy1*-YFP16 mice; “cKO”, experimental group with ChAT-Cre^mut/wt^ × *Atg7*^fl/fl^ × *Thy1*-mito-CFP × *Thy1*-YFP16; n ≥ 19 axons from 3 mice per group). **g**, Reactive oxygen species of presynaptic mitochondria (mito-ROS) in acute nerve-muscle explants. Mito-Grx1-roGFP2 signal ratio at 405/488 nm excitation (“CTR”, ChAT-Cre^wt/wt^ × *Atg7*^fl/fl^ × *Thy1*-mito-Grx1-roGFP2 pooled with ChAT-Cre^mut/wt^ ^or^ ^wt/wt^ × *Atg7*^wt/wt^ × *Thy1*-mito-Grx1-roGFP2 mice; “cKO”, experimental group with ChAT-Cre^mut/wt^ × *Atg7*^fl/fl^ × *Thy1*-mito-Grx1-roGFP2; n ≥ 16 axons from 4 mice per group). **h–m**, Distal degeneration and swelling of motor axon terminals in triangularis sterni muscles of 11– 20-week-old mice (Left column: “CTR”, ChAT-Cre^wt/wt^ × *Atg7*^fl/fl^ × *Thy1-*YFP16 pooled with ChAT-Cre^mut/wt^ ^or^ ^wt/wt^ × *Atg7*^wt/wt^ × *Thy1-*YFP16 mice; Right column: “cKO”, ChAT-Cre^mut/wt^ × *Atg7*^fl/fl^ × *Thy1-*YFP16; anti-Caspr-antibody, yellow; α-bungarotoxin, magenta; YFP16, white). **h**,**i**, Confocal projections of intercostal nerve fascicles (proximal, ‘stem’ axons), **j**,**k**, terminal motor axon branches (boxes indicate positions of the images underneath this row with magnified views, enumerating the internodes of selected axons from distal-most to proximal). **l**, Axon calibers of internodes, quantified from distal-most (1°) to more proximal (4°) (“Atg7 CTR”, ChAT-Cre^wt/wt^ × *Atg7*^fl/fl^ × *Thy1*-mito-Dendra pooled with ChAT-Cre^mut/wt^ ^or^ ^wt/wt^ × *Atg7*^wt/wt^ × *Thy1*-mito-Dendra mice; “Atg7-cKO”, experimental group with ChAT-Cre^mut/wt^ × *Atg7*^fl/fl^ × *Thy1*-mito-Dendra littermates; n ≥ 70 axons from 3 mice per group). **m**, Denervation of neuromuscular endplates (Top arrow: axon completely vacated the postsynaptic territory; Bottom arrow: partially denervated junction). Scale bars in **a**,**c** 10 µm (top), 2 µm (magnified views, bottom); **b**,**d** 300 nm; **h**,**i** 10 µm; **j**,**k**,**m** 20 µm. Violin plots in **e** show data distribution with median. Box plots in **f**,**g**: median, quartile 1 - quartile 3, whiskers 95th percentiles. Mann-Whitney test was used to determine significance in **e**,**f**,**g**,**l**: ∗∗∗p < 0.001; ∗p < 0.05; n.s. non-significant.

Interestingly, however, spherical mitochondria with damaged ultrastructure (**Fig. 4a**–**d**), low membrane potential (**Fig. 4f**), and high mitochondrial reactive oxygen species (ROS) levels (**Fig. 4g**) accumulated in junctions of *Atg7*-cKO mice. The axon terminals filled with this debris were abnormally swollen and degenerated in *Atg7*-cKO mice (**Fig. 4a–d,j,k,m**). While generic defects in synaptic structure and function were described in *Atg7*-cKO mice (Rudnick et al., 2017), we observed a striking spatial distribution of the initial phenotype with distal-to-proximal tapering of swellings, being most pronounced at NMJs and the distal internodes, but absent more proximally (**Fig. 4j**–**l**; see also Friedman et al., 2012; Komatsu et al., 2007 for other neuron types), but being not apparent at more proximal internodes and stem axons (**Fig. 4h,i**). The distally pronounced accumulation of dysfunctional organelles at NMJs suggested to us that damaged mitochondria are indeed removed from junctions by *Atg7*-related autophagy in large numbers, consistent with our direct observations of capture at the exit point. However, the transport results also suggest that ‘capture’ is rather an upstream event that occurs prior to Atg7-mediated degradation, preventing disrupted mitochondria from leaving the junction, even in *Atg7*-cKO mice. This seems to be in line with retrograde (‘passing’) transport, but not anterograde flux, being reduced when compared to controls with wildtype Atg7 expression (**Fig. 4e**), as would be predicted if damaged mitochondria would still be captured, but could not be degraded and as a result accumulated over time (see **Fig. 2**).

### Capture of mitochondria depends on optineurin but does not require the PINK1/parkin pathway

Since our results indicated a selective mechanism of recognizing damaged mitochondria for synaptic degradation, we examined whether well-established mitophagy pathways are involved. We first tested whether PINK1 and parkin, two proteins that interact to label mitochondria with polyubiquitin chains for the initiation of mitophagy (Geisler et al., 2010; Matsuda et al., 2010; Narendra et al., 2008, 2010; Vives-Bauza et al., 2010), influence the capture mechanism at synaptic terminals. Both proteins detach damaged mitochondria from the transport machinery to promote local axonal mitophagy *in vitro* (Ashrafi et al., 2014; Wang et al., 2011). Notably, we observed patchy degeneration of axon terminals in adult *Pink1-Prkn*-double-knock out (DKO) mice, which went along with reduced retrograde transport in the stem axon (**Extended Data Fig. 5a–e**; Rogers et al., 2017). As these adult NMJs, however, showed clear signs of sprouting and other morphological changes, we concluded that a proper assessment of mitochondrial mass turnover would not be possible at this stage. We therefore performed mitochondrial capture experiment in *Pink1-Prkn*-DKO axon terminals at an earlier timepoint (3 weeks of age), before the onset of NMJ disruptions and the corresponding transport phenotype (**Extended Data Fig. 5f**, Mann-Whitney-tests for mitochondrial volume flux: Anterograde, p ≥ 0.05; Retrograde p ≥ 0.05). In this setting, surprisingly, absolute volume flux rates in *Pink1*-KO (**Fig. 5a**) and *Pink1-Prkn*-DKO were unchanged (**Fig. 5c**), and capture rates were not decreased compared to littermate controls as would be expected if capture required the PINK1/parkin mechanism (**Fig. 5b,d**; Ctr: 54% capture, DKO: 62% capture; per group ≥ 29 tracked mitochondria during 20 ≥ ‘NMJ hours’, χ²-test p ≥ 0.05). Overall, this suggests that a different (PINK1/parkin-independent) mechanism can retain damaged mitochondria in exit points, putatively with the involvement of other ubiquitinylation-modifying enzymes (such as MITOL and others; Evans & Holzbaur, 2019b; Villa et al., 2018; Yonashiro et al., 2006). In favor of this possibility, we found that immunostainings against ubiquitin showed elevated labeling levels at synaptic exit points compared to other areas of the terminal (**Fig. 5e–h**).

**Fig. 5:**
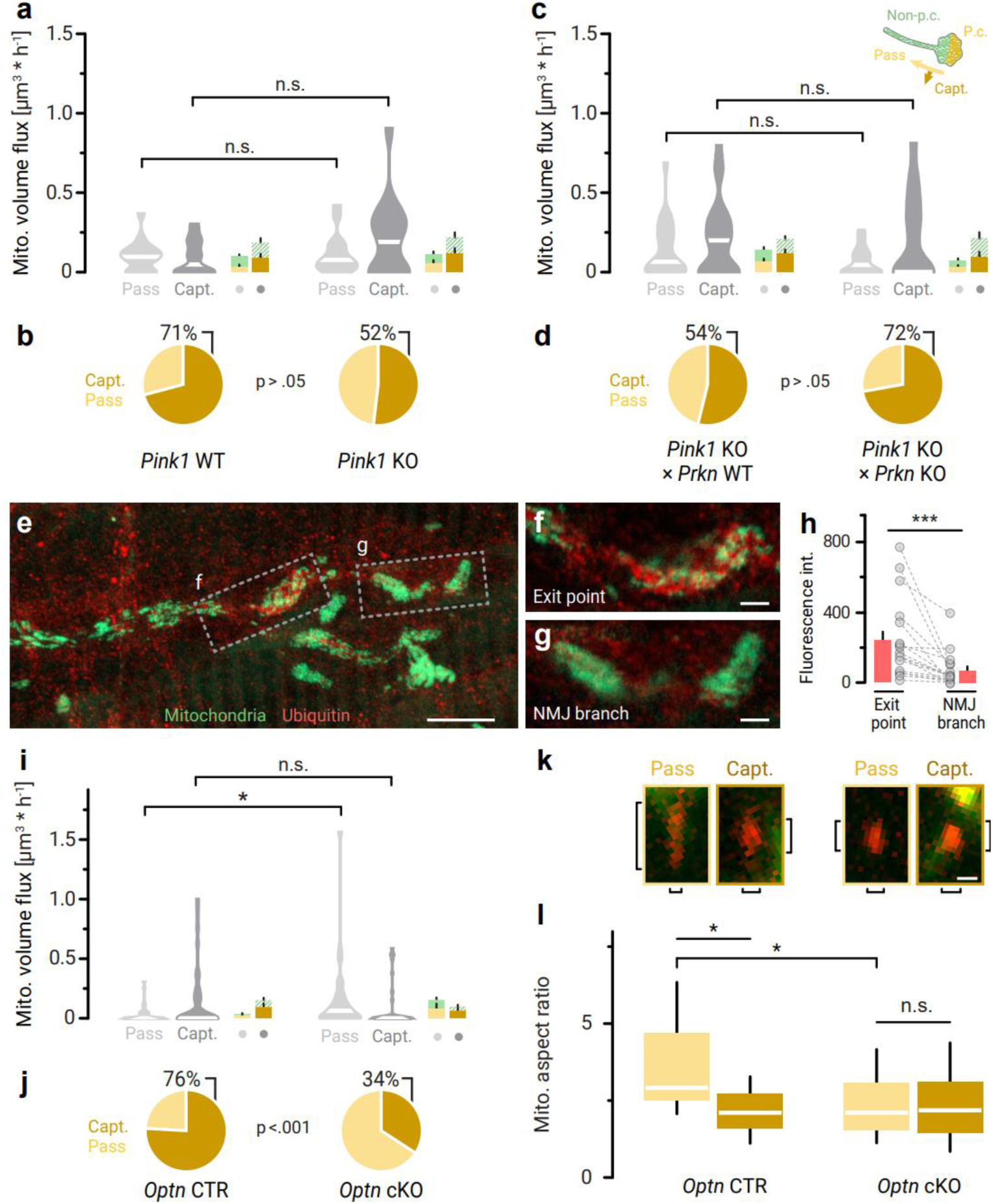
Mitochondrial ‘capture’ in motor axons depends on optineurin but does not require the PINK1/parkin pathway. **a**,**b**, In motor axon terminals of three-week old *Pink1* gene-deleted mice, mitochondrial capture is not significantly altered (Left block: control group of *Pink1*^+/+^ (wildtype, WT) × *Thy1*-mito-Dendra mice; Right: experimental group with *Pink1*^-/-^ (knockout, KO) × *Thy1*-mito-Dendra littermates; n ≥ 8 mice per genotype, ≥ 15 ‘NMJ imaging hours’). **a**, Mitochondrial volume flux at synaptic exit points (Legend: as in Fig. 2c) (n ≥ 15 axons per genotype). **b**, Fractions of p.c. retrogradely moving mitochondria that were captured at synaptic exit points (n ≥ 24 mitochondria per genotype). **c**,**d**, A combined gene deletion of Pink1 and parkin does not affect mitochondrial capture (Left: control group of three-week old *Pink1*^-/-^ (KO) × *Prkn*^wt/wt^ × *Thy1*-mito-Dendra mice; Right: experimental group with *Pink1*^-/-^ (KO) × *Prkn*^ko/ko^ × *Thy1*-mito-Dendra littermates; n ≥ 10 mice per genotype, ≥ 15 ‘NMJ hours’). **c**, Mitochondrial volume flux at synaptic exit points (Legend: as in Fig. 2c) (n ≥ 15 axons per genotype). **d**, Fractions of p.c. retrogradely moving mitochondria that were captured at synaptic exit points (n ≥ 18 mitochondria per genotype). **e**–**h**, Accumulation of ubiquitin in synaptic exit points. **e**, Confocal maximum projection of an antibody staining against ubiquitin (red) in a motor axon terminal with Dendra-labelled mitochondria (green). Boxes outline the location of the single optical sections shown in **f**, synaptic exit point, and **g**, NMJ branch tip. **h**, Anti-ubiquitin content (mean intensity pixel values) in synaptic exit points and distal NMJ branches (n ≥ 18 axons from 2 *Thy1*-mito-Dendra mice). **i**,**j**, A cholinergic-neuron specific gene deletion of optineurin allows mitochondria to ‘pass’ through the synaptic exit point at a greater rate (Left block ‘CTR’: control group of ChAT-Cre^wt/wt^ × *Optn*^fl/fl^ × *Thy1*-mito-Dendra mice; Right ‘cKO’: experimental group with ChAT-Cre^mut/wt^ × *Optn*^fl/fl^ × *Thy1*-mito-Dendra littermates; n ≥ 18 mice per genotype). **i**, Mitochondrial volume flux at synaptic exit points (Legend: as in Fig. 2c) (n ≥ 29 axons per genotype). **j**, Fractions of p.c. retrogradely moving mitochondria that were captured at synaptic exit points (n ≥ 29 mitochondria per genotype). **k,l**, Morphological properties of ‘captured’ and ‘passing’ mitochondria in Optn-cKO and control mice (analyzed from the movies in **i,j**); **k** shows retrogradely moving p.c. mitochondria (red) in the synaptic exit point (left pair: Optn control, right pair: Optn-cKO). Vertical brackets represent mitochondrial length, horizontal brackets mitochondrial width, to illustrate how measurements in **l** were taken. **l**, Mitochondrial aspect ratio, which denotes the ratio of mitochondrial length and width, for retrogradely moving p.c. mitochondria from each genotype and class (n ≥ 13 mitochondria per group). Scale bars in **e** 10 µm; **f**,**g** 2 µm; **k** 500 nm for all image panels. Violin plots in **a,c,i** show data distribution and median values. Bar graphs in **h**: mean ± s.e.m.; pairings between individual datapoints are indicated by dotted lines. Box plots in **l**: median, quartile 1 - quartile 3, whiskers 95^th^ percentiles. Mann-Whitney test was used to determine significance in **a**,**c**,**i**, a Wilcoxon (paired) test was used in **h**, a Kruskal Wallis test (p < 0.01) with multiple comparisons correction was used to determine significance in **l**: ∗∗∗p < 0.001; ∗p < 0.05; n.s. non-significant; a χ²-test was used in pie charts (**b**, p = 0.1658; **d**, p = 0.1887; **j**, p = 0.0006).

In addition to the ubiquitination system per se, mitophagy also depends on specific adaptor proteins that link the ubiquitin-tagged mitochondrion to autophagosomal proteins, such as LC3 (Stavoe & Holzbaur, 2019). Notably, some of the adaptors, specifically optineurin (OPTN), are linked to ALS (Markovinovic et al., 2017), which suggests a specific involvement in mitophagy in motor neurons. Indeed, when we deleted optineurin selectively in cholinergic motor axons (ChAT-Cre^mut/wt^ × *Optn*^fl/fl^, abbreviated below to *Optn*-cKO), we found that capture rates were reduced by more than half, from 76% in the control to 34% in the *Optn*-cKO (**Fig. 5j**, p < 0.01, χ²-test; n ≥ 38 tracked mitochondria per group; ChAT-Cre^mut/wt^ × *Optn*^fl/fl^ mice compared to ChAT-Cre^wt/wt^ × *Optn*^fl/fl^ littermate controls). Mitochondrial volume flux also increased 4-fold in the passing population (Controls: 0.04 ± 0.01 µm³/h, cKO: 0.16 ± 0.06 µm³/h, p < 0.05, Mann-Whitney-U-test), suggesting that when lacking optineurin, mitochondria that would otherwise be captured in exit points, are now able to pass into the axon (**Fig. 5i**). In line with this interpretation, passing mitochondria in the *Optn*-KO axons were rounder in shape than in controls, resembling their normally captured counterparts (**Fig. 5k,l**; shape factors ‘passing’ mitochondria: Ctr 3.5 ± 0.4, cKO 2.3 ± 0.2; ‘Captured’ mitochondria: Ctr 2.2 ± 0.1, cKO 2.3 ± 0.3). Retrograde mitochondrial volume flux was not significantly altered in intercostal nerve stem axons (**Extended Data Fig. 5g**) (Ctr: 0.15 ± 0.02 µm³/min, KO: 0.17 ± 0.04 µm³/min, p > 0.05, Mann-Whitney-U-test), indicating that optineurin is required predominantly in synaptic terminals for efficient mitochondrial capture. Together, these data reveal a non-canonical mitophagy mechanism that degrades most of the synaptic mitochondrial volume that is removed from motor axons.

## DISCUSSION

Mitostasis—the long-term maintenance of healthy mitochondria in all cell compartments—taxes neuronal cell biology and remains enigmatic (Misgeld & Schwarz, 2017; Stavoe & Holzbaur, 2019; Pekkurnaz & Wang, 2022; Hill & Colon-Ramos, 2020; Coleman & Hoke, 2020). Our quantitative exploration of mitochondrial turnover in motor axons, which are amongst the most extensive and disease-prone projections in the mammalian nervous system, now resolves several seminal questions, namely the location of the major site of synaptic mitophagy, as well as the key underlying cell biological and molecular mechanisms. The distal mechanism of neuronal mitophagy that we propose combines features of several previous descriptions (e.g., Maday, 2016; Ashrafi et al., 2014; McWilliams et al., 2018; Han et al., 2020; Lie et al., 2021; Sheng & Cai, 2012; Sheng, 2014), but overall amounts to a distinct model of how large projection neurons maintain their most distal mitochondrial outposts. In a nutshell, our model proposes that in NMJs, functionally impaired mitochondria or their fragments initiate a form of local retrograde transport and are then sorted out as they exit the synapse or traverse distal nodes (**Fig. 1 and 2**). Here, they are rerouted to a degradation ‘hotspot’ while being decorated with ubiquitin, engulfed by autophagic membranes to finally merge with local lysosomes (**Fig. 3**). The transition of retrograde transport into mitophagy involves an optineurin-mediated but PINK1/parkin-independent ‘capture’ mechanism (**Fig. 5**) that is over-active early in models of motor neuron disease (**Fig. 2**).

Together, this illuminates several key aspects of mitochondrial quality control in motor neurons:

(1) *Most mitophagy in motor axons is synaptic but still linked to transport*: Our work argues against the notion that mitochondrial degradation in neurons is mostly somatic and perhaps even the raison d’etre of retrograde transport (Sheng & Cai, 2012; Sheng, 2014). Moreover, we also refute that axons cannot contain sufficiently mature lysosomes or endolysosomes to execute classical degradative functions (Han et al., 2020; Lie et al., 2021). In converse, our results are in line with previous data showing that autophagic degradation of mitochondria involves initiation of retrograde motility (Maday, 2016; Neisch et al., 2017), which, however, ceases beyond a certain point of mitochondrial dysfunction as transport adaptors are degraded (Ashrafi et al., 2014; Wang et al., 2011). While our data clearly establishes this mechanism for motor axons, whether distal degradation in specific axon locations is a general feature of all neurons remains open—while motor axons are important and disease-relevant, their large terminal synapses and thick Schwann cell-myelinated axons are not typical of most CNS axons, which are thin and carry en passant synapses. However, autophagy has been described as an important mechanism for CNS synapse function and development (Stavoe & Holzbaur, 2019), suggesting that the required degradative machinery is also present in central axons. Moreover, the reported accumulation of mitochondrial material at axonal branch points (which in myelinated axons are also Nodes of Ranvier and sites of synapses in the CNS) might hint at the location of mitophagic hot spots in axons with an en passant synaptic arrangement (Courchet et al., 2013; Plucinska et al., 2012; Spillane et al., 2013). Either way, given the unique vulnerability of motor neurons to specific defects in mitophagy-related pathways, an NMJ-specific mechanism of mitophagy is of substantial biomedical significance, especially as such a peripheral site of substantial mitophagy has unique pharmacological accessibility.

(2) *Mitophagy is clustered perinodally*: The fact that perinodal sites are hotspots for mitochondrial degradation strengthens previous notions that paranodal ‘Schwann cell network’ around peripheral nodes of Ranvier can mediate degradative and detoxifying processes (Gatzinsky et al., 1997). While previous models of glial ‘transmitophagy’ (Davis et al., 2014; Burdett and Freeman, 2014; Phinney et al., 2015) suggest eventual transfer of degrading mitochondria to the glial surround, our observations of mature motor axons and NMJs showed no evidence of this, even though at developing NMJs, transfer of axonal material to glia is readily observable (Bishop et al., 2004). While we cannot definitively prove that nodes are necessary for the capture mechanism (as most models lacking myelination or Schwann cells have additional axonal phenotypes; Djannatian et al., 2019) or explain the reason for this intriguing location, it is worth noting that paranodal areas contain a specialized actin cytoskeleton. As optineurin has a myosin-binding domain (Ryan & Tumbarello, 2018; Slowicka et al., 2016), this might mediate the observed mitochondrial capture events. However, whether such cytoskeletal specializations are also present on the distal side of the last heminode—the site of the most effective retrograde filter—is currently not known.

(3) *Decentralized mitophagy allows scaling with motor unit size*: Irrespective of mechanisms, the extent of mitochondrial degradation in the periphery of motor axons that our work reveals is staggering: Each terminal axon branch contains an array of sequentially arranged paranodal checkpoints, establishing a semi-autonomous system that accounts for at least 75% of presynaptic mitochondrial turn-over (**Fig. 1**). The synaptic mitochondrial population in a motor neuron is probably orders of magnitude bigger than somatic content (which, e.g. for layer 5 ‘upper’ motor neurons is ∼300; Lin et al., 2019; with probably a somewhat larger pool for the bigger ‘lower’ motor neurons that we study; Tamada, 2023). Based on an estimate of ∼50 NMJs in a small motor unit as found in thin muscles like the triangularis sterni (Keller-Peck et al., 2001; Lu et al., 2009) and around 300 to 500 mitochondria in a typical NMJ (based on our own ultrastructural reconstructions), we estimate the number of a small motor neuron’s synaptic mitochondria to exceed 15000 (Misgeld and Schwarz, 2017). Given an anterograde delivery rate of 4 mitochondria per hour (**Fig. 1**; Misgeld et al., 2007), we can extrapolate that the presynaptic pool is turned over within 3 to 5 days, which seems plausible considering previously reported mitochondrial protein half-lives (Dorrbaum et al., 2018). Here, the advantage of a decentralized local degradation system becomes obvious: It is inherently scaleable since its capacity relies directly on the number of axon terminals, precluding the risk that the stem axon or the soma get overwhelmed by an excessive stream of retrogradely moving mitochondrial debris. In addition, such a system is inherently independent of the distance to the soma and can adapt to changes in geometry that physiologically result from remodeling and plasticity, both in development and in aging (Sanes & Lichtman, 1999).

(4) *The non-canonical molecular machinery might link to specific neurodegenerative phenotypes*: Beyond these basic physiological insights, our findings also have significance in the context of disease. It appears from our work that the quantitatively most important mitophagic system in motor axons employs optineurin, but PINK1 and parkin are not absolutely required. This suggests the involvement of a parkin-independent form of mitophagy, several of which have been described, including ones that involve NIX, FKBP4, or other mitophagy receptors (Teresak et al., 2022). However, most of these are ubiquitin-independent and can directly link to the phagophore membrane and do not involve optineurin – leaving the exact molecular pathway involved in mitophagy at NMJs unresolved, but rather suggesting an alternative ubiquitylation pathway (Evans & Holzbaur, 2019b; Villa et al., 2018; Yonashiro et al., 2006). Still, the involvement of optineurin, but not PINK1/parkin, could illuminate the cell type- and region-specificity of synapse damage observed in neurodegenerative diseases that are linked to mitochondrial dysfunction. This certainly chimes with the manifestation of human optineurin mutations as ALS (Cirulli et al., 2015; Maruyama et al., 2010) and not PD, which is linked to PINK1 and parkin (Pickrell & Youle, 2015). Our data furthermore imply a mechanistic entanglement of defects in retrograde mitochondrial transport and degradation, which are often observed concomitantly in neurodegenerative diseases (Abeliovich & Gitler, 2016; Evans & Holzbaur, 2019a; De Vos et al., 2017; Killackey et al., 2020). Disruption of short-range mitochondrial transport within axon terminals would decrease a neuron’s capacity for peripheral degradation of damaged mitochondria, while defective bulk autophagy would secondarily alter long-range axonal transport towards the soma.

There are also two unresolved aspects of our model regarding mitochondrial trafficking to be clarified in future work: The origin of the captured mitochondria and the fate of the passing ones. For the origin of the captured mitochondria, two not necessarily exclusive possibilities are likely. First, the retrogradely traveling organelles could be individual synaptic mitochondria that are fated for degradation after having reached the end of their utility. Indeed, our photo-tagging experiments argue for the existence of stably distinct organelles in the synapse rather than a mitochondrial network that is dynamically interconnected by fusion and fission as typically found in cultured cells, as in such a network a photo-tagged subpopulation would quickly be dissipated. Moreover, the apparent volume of the captured mitochondria was not clearly distinct from the mostly stable mitochondria that electron microscopy revealed in the synapse (cf. **Fig. 1g** and **Fig. 3**; which show a mean volume of mitochondria of 0.09 ± 0.01 µm^3^ for captured and 0.06 ± 0.01 µm^3^ for passed mitochondria vs. a mean volume of resident mitos in EM reconstructions of 0.05 ± 0.03 µm^3^ corrected for 30% tissue shrinkage). This further supports the notion that the transported organelles could be ‘complete’ mitochondria. However, we can also not exclude the possibility that the cleared material is the product of local and restricted dynamics involving mitochondrial fusions, followed by sorting of material and finally fission, likely of the type described as asymmetric and degradative in recent *in vitro* work (Kleele et al., 2021). This notion might be supported by the fact that in shape, albeit not in volume, the captured mitochondria are more spherical than anchored or long-distance traveling mitochondria.

Notable in this context is the observation that the passing mitochondria, which enter the axon from the synapse, do not match in size those that travel retrogradely in the stem axon (**Fig. 1**). Based on our mitoKeima experiments, these are not mitolysosomes, but rather *bona fide* mitochondria. This either implies that the retrogradely-traveling mitochondria originate from a different source, e.g. within the axon itself rather than the synapse, and that the repetitive perinodal filters eventually capture most if not all mitochondria that have left the synapse. Or, alternatively, en passant fission and fusion events in the axon could reshape this organelle pool—but this seems unlikely, given that such mitochondrial dynamics are not really observed in mature motor axons. Ultimately, we must concede that not only the source, but also the purpose of retrograde transport of mitochondria is increasingly opaque. The latter could include i.a.: (*i*) retrograde shuttling of mitochondria from the proximal axon before, while or after they are being packaged in mitophagy- or bulk autophagy-related membranes (Maday et al., 2012), destined for somatic mitophagy (Miller and Sheetz, 2004; Ashrafi et al., 2014; Lin et al., 2017), as proximal perinodal sites in stem axons do not seem to have degradative capacity (see Results); (*ii*) return for repair or just general admixing of mitochondrial content to avoid local build-up of damaged mitochondrial material (Wong et al., 2012; Mandal et al., 2021), which indeed can accumulate in neurons in lysosomal storage disorders (Rowan & Lake, 1995); (*iii*) retrograde signaling to the soma, putatively about the size of the peripheral mitochondrial pool, as has been suggested for other cargoes (Rishal & Fainzilber, 2014). We did not specifically address the identity of this ‘passed’ population in our work. Still, we think a plausible model would be to assume a relatively contiguous ‘baseline’ stream of organelles (including mitochondria) derived from the axon proper that is moving toward the soma destined for or even in the process of bulk autophagy. This long-distance process would be augmented by the local degradative mechanisms in the neuronal periphery described here. This system is more attuned to both neuronal geometry and acute changes in mitochondrial health—and even in small motor units like the one studied here, easily exceeds the central mechanism in terms of the degraded volume. Thus, employing various mechanisms of mitochondrial quality control may serve as a means of scaling but also as a backup mechanism in situations of distress or disruption of certain pathways—which could explain that any single disruption of mitostasis mechanisms will typically only manifest as a slowly progressive problem.

In summary, the peripheral mitostasis mechanism that we describe here can elucidate two of the puzzling characteristics of mitochondrially-driven neurodegeneration: the cell-type specificity, as well as the late onset “dying-back” pattern of many of such pathologies (Coleman & Hoke, 2020). Whether and how such insights could be leveraged to intervene in such ‘mitostasis disorders’ can now be probed in future work.

## ACKNOWLEDGEMENTS

We are grateful to M. and N. Budak, as well as M. Korica for excellent animal husbandry and K. Kellermann for veterinary support. We thank E. Bader, E. Cetin, O. Distl, Y. Hufnagel, J.-N. Dreessen, R. Karl, K. Karg, N. Sulzer, M Schetterer, and K. Wullimann for their help in collecting data, as well as outstanding technical and administrative support. We also thank the Lichtman and Sanes labs (Harvard University) for their generous provision of *Thy1*-reporter mice and coated Kapton tap; D. Edbauer, Q. Zhou, and W. Wurst (all DZNE) for the provision of the GA-CFP and the Pink1- and Prkn-KO mice, respectively. We thank A. Stout and the University of Pennsylvania, Cell and Developmental Biology Microscopy Core facility for help with Airyscan imaging. We thank L. Godinho for valuable input on a previous version of this manuscript.

Work in T.M.’s lab is supported by the Deutsche Forschungsgemeinschaft (Mi 694/8-1 “In vivo analysis of mitochondrial dynamics and fate in axons and synapses”, as well as Mi 694/7-1, Mi 694/9-1 A03-ID 428663564, FOR Immunostroke and via TRR 274/1 2020, project C02 – ID 408885537) and the ERC under the European Union’s Seventh Framework Program (FP/2007-2013; ERC Grant Agreement n. 616791). T.M. and M.S. are members of and supported by the German Center for Neurodegenerative Diseases (DZNE), TRR 274/1, project Z01, and a Chan Zuckerberg Initiative “Advancing Imaging Through Collaborative Projects” grant. High-resolution microscopy was supported via a DFG instrumentation grant (INST95/1755-1 FUGG, ID 518284373). T.M., M.S. and M.S.B. receive support from the Munich Center for Systems Neurology (SyNergy EXC 2145; Project ID 390857198); M.S.B. is also supported by the DFG (project ID: 450131873), and the DGM foundation (ID Le3/1). T.K. receives grant support from the Swiss National Science Foundation (SNSF, 310030_215716); S.E: was supported by the DFG (project ID 403584255–TRR267) and the Federal Ministry of Education and Research (BMBF) in the framework of the Cluster4future program (CNATM - Cluster for Nucleic Acid Therapeutics Munich). M.L. was awarded a Hans-Fischer Fellowship for this project by the TUM Institute for Advanced Study (TUM-IAS), which was hosted by T.M. and supported N.M.’s salary. N.M. was further supported by the Graduate School of TUM (TUM-GS) and received a fellowship from the TUM School of Medicine. A.I. was a student of the MSc program ‘Biomedical Neuroscience’ and received support from the Elite Network Bavaria (ENB).

## AUTHOR CONTRIBUTIONS

N.M., T.K., M.S.B., and T.M. conceptualized the project and experiments. T.M. and M.B. administered the project and the mouse lines. They also provided supervision to N.M. and B.P.; N.M. supervised A.I. N.M. performed most of the investigation, including ex vivo imaging, structural microscopy (with M.L.’s support for Airyscan imaging), and motor neuron disease modeling experiments (with M.S.B. help in managing mouse lines), with contributions from T.K., B.P. and A.I. for pulse-chase imaging, confocal microscopy and data analysis. T.M., M.L., B.P., and M.B. established and characterized mouse lines. M.S. prepared and imaged samples for electron microscopy in collaboration with N.M., N.M. analyzed and provided visualizations for rendered electron micrographs. P.A. and S.E. designed and generated AAV vectors and cloned constructs. N.M. coordinated and contributed analysis of data across all experimental approaches and designed the final data representation with input from T.M.. N.M. and T.M. wrote the original draft of the paper with input from all authors, who also reviewed and edited the manuscript. S.E., M.B., and T.M. were responsible for funding acquisition and provided instrumentation resources.

## EXTENDED DATA FIGURE LEGENDS

**Extended Data Fig. 1:**
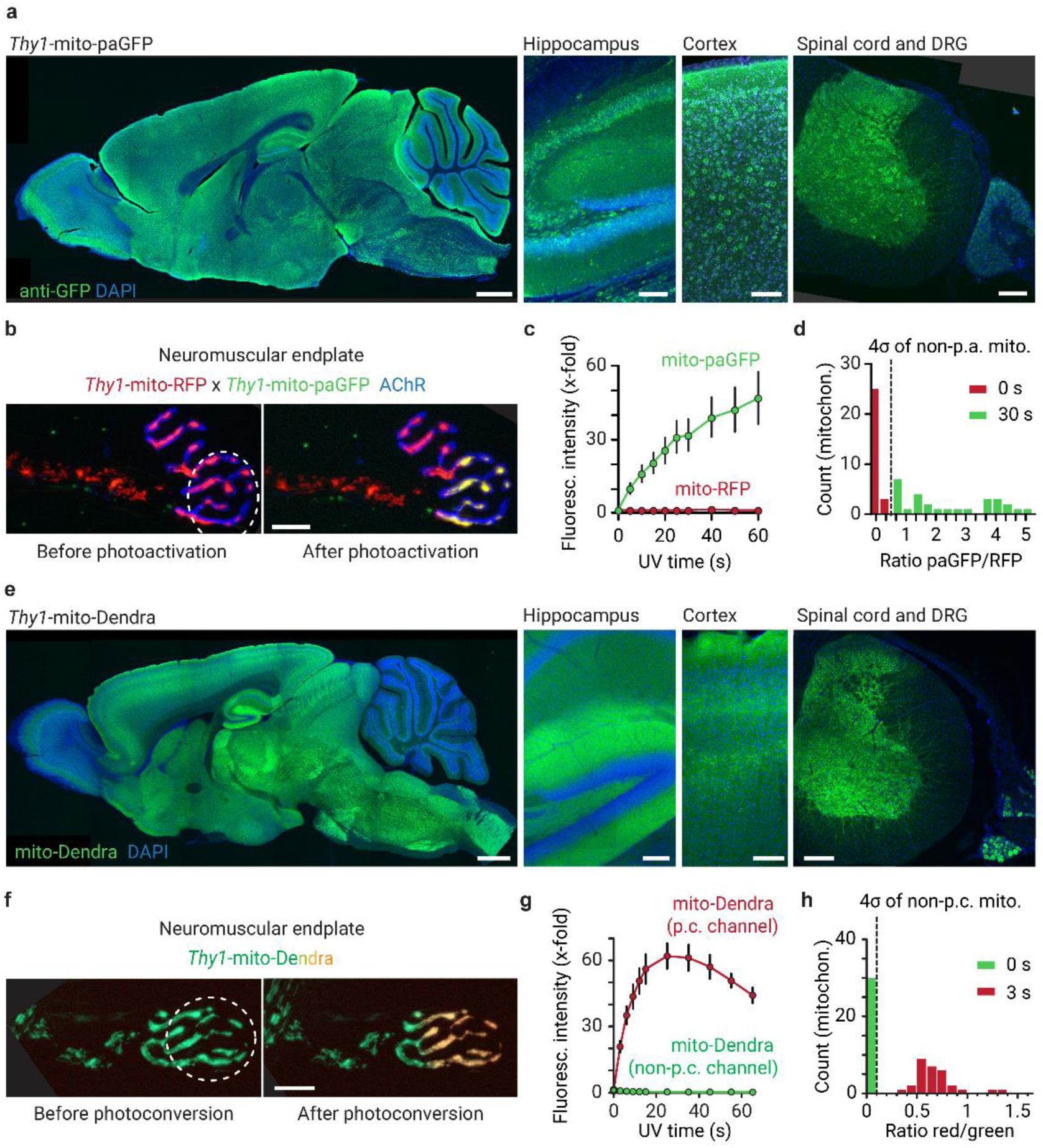
Expression pattern and photoactivation / -conversion in newly generated *Thy1*-mito-paGFP and *Thy1*-mito-Dendra transgenic mouse lines. **a**, The mito-paGFP transgene is widely expressed throughout the central and peripheral nervous system (stained with anti-GFP-antibody). Left to right: sagittal brain section, hippocampus, cerebral cortex, horizontal section of spinal cord and a dorsal root ganglion (DRG). **b**, Confocal projections of motor axon terminals before and after photoactivation with 290 µW 405 nm light in the circled region for 30 s (acute nerve-muscle explant, *Thy1*-mito-paGFP × *Thy1*-mito-RFP; postsynaptic acetylcholine receptors stained with α-bungarotoxin, blue). **c**, Photoactivation (“p.a.”) efficiency with ascending doses of 405 nm illumination. 30 s illumination yielded ∼30fold increase in paGFP fluorescence intensity (∼8.7 mW*s; n = 12 NMJs from 4 *Thy1*-mito-paGFP × *Thy1*-mito-RFP mice). **d**, Distribution of paGFP/RFP fluorescence intensity ratio before and after p.a. (settings as in **b**; n = 28 mitochondria from the experiments in **c**). Dashed line indicates the 4× standard deviation distance from the mean of the non-p.a. mitochondria population. **e**, Expression of the mito-Dendra transgene throughout the nervous system (endogenous non-photoconverted Dendra fluorescence). Left to right: as in **a**. **f**, Widefield projections of motor axon terminals before and after photoconversion with 290 µW 405 nm illumination in the circled region for 3 s (from green to red fluorescence; acute nerve-muscle explant, *Thy1*-mito-Dendra). **g**, Photoconversion (p.c.) efficiency with 405 nm light. 3 s illumination yielded ∼20fold increase of Dendra-red (p.c.) brightness, corresponding to ∼30fold increase in Dendra-red/green ratio (∼870 µW*s; n = 10 NMJs, 2 *Thy1*-mito-Dendra mice). **h**, Distribution of Dendra-red/green ratio before and after p.c. (as in **f**; n = 30 mitochondria from the experiments in **g**). Dashed line: as in **d**. Scale bars in **a** and **e** (from left to right): 1 mm, 50 µm, 50 µm, 200 μm; in **b** and **f** 10 μm. **c** and **g** show mean ± s.e.m.

**Extended Data Fig. 2:**
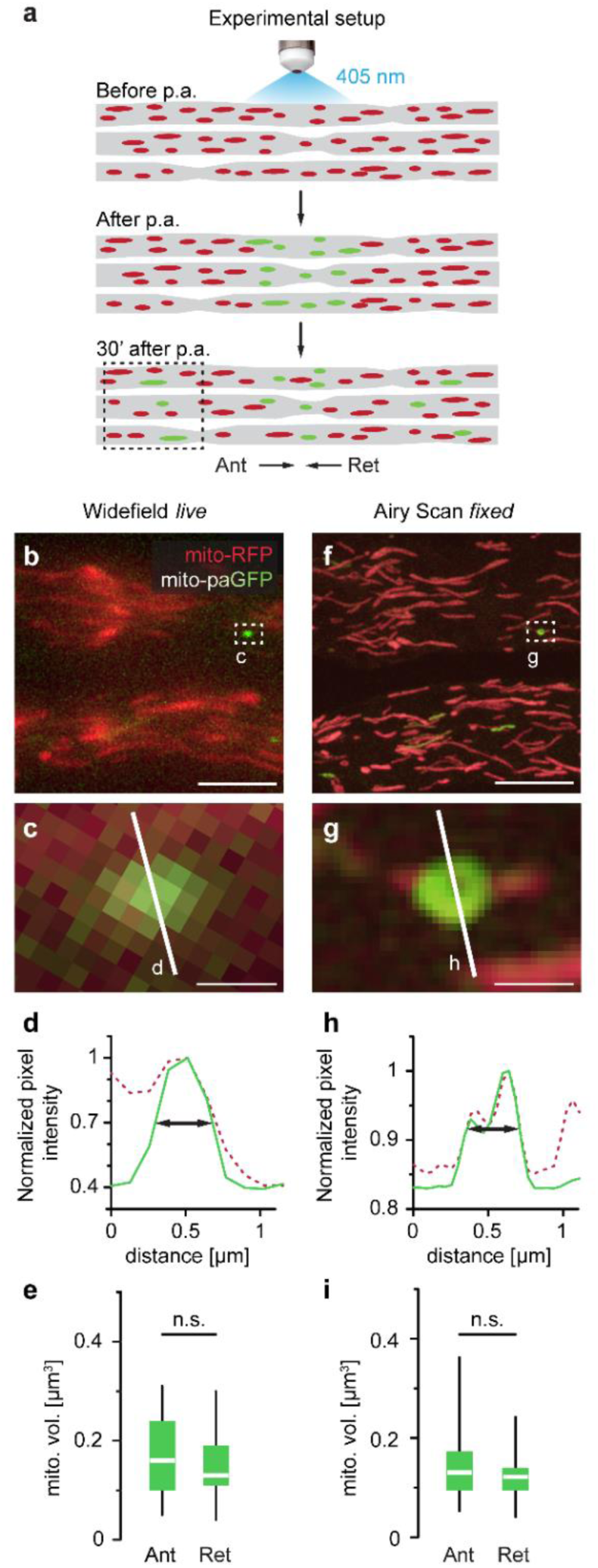
High-resolution volume measurements of mitochondria undergoing axonal transport. **a**, Experimental setup: Neuronal mitochondria containing both (non-fluorescent) photoactivatable GFP (paGFP; Patterson & Lippincott-Schwartz, 2002) and (fluorescent) RFP were illuminated with 405 nm light in a 200-µm stretch of intercostal nerve, thus photoactivating paGFP (green) and marking the mitochondria for later observation (acute nerve-muscle explants from *Thy1*-mito-paGFP × *Thy1*-mito-RFP mice; Breckwoldt et al., 2014). 30 min after photoactivation (p.a.), individual green-fluorescent mitochondria undergoing axonal transport were identified in regions adjacent to the photoactivation site (**b**,**c**). Following fixation (which paGFP withstands), displaced mitochondria were imaged using the higher resolution of Airy Scan microscopy (**f**,**g**). **b**, Widefield live image adjacent to the photoactivation site. **c**, A single photoactivated mitochondrion (position is outlined by the box in **b**). **d**, Intensity profile of pixels across the mitochondrion in **c** (pixel locations indicated by the line in **c**; green: pa-GFP signal, red: RFP signal). Arrows indicate the width of the paGFP-intensity curve at half of its amplitude. **e**, Mitochondrial volume of photoactivated mitochondria displaced in anterograde and retrograde directions, measured in widefield images (n ≥ 15 axons, ≥ 4 mice per group). **f**, Overview of the region shown in **b** after sample fixation and re-location. **g**, Airy Scan image of the photoactivated mitochondrion shown in **c**; position of image outlined by box in **f**. **h**, Intensity profile of pixels along the line shown in **g**. Colors and symbols as in **d**. **i**, Mitochondrial volume of photoactivated mitochondria displaced in anterograde and retrograde directions, measured in Airy scan image stacks (for calculation, see Methods; n ≥ 18 axons, ≥ 3 mice per group). Scale bars in **b**,**f** 5 μm, in **c**,**g** 500 nm. Box plots: median, quartile 1 - quartile 3, whiskers 95th percentiles. Unpaired *t*-test was used to determine significance in **e** and Mann-Whitney test in **i**: n.s.: non-significant, p ≥ 0.05).

**Extended Data Fig. 3:**
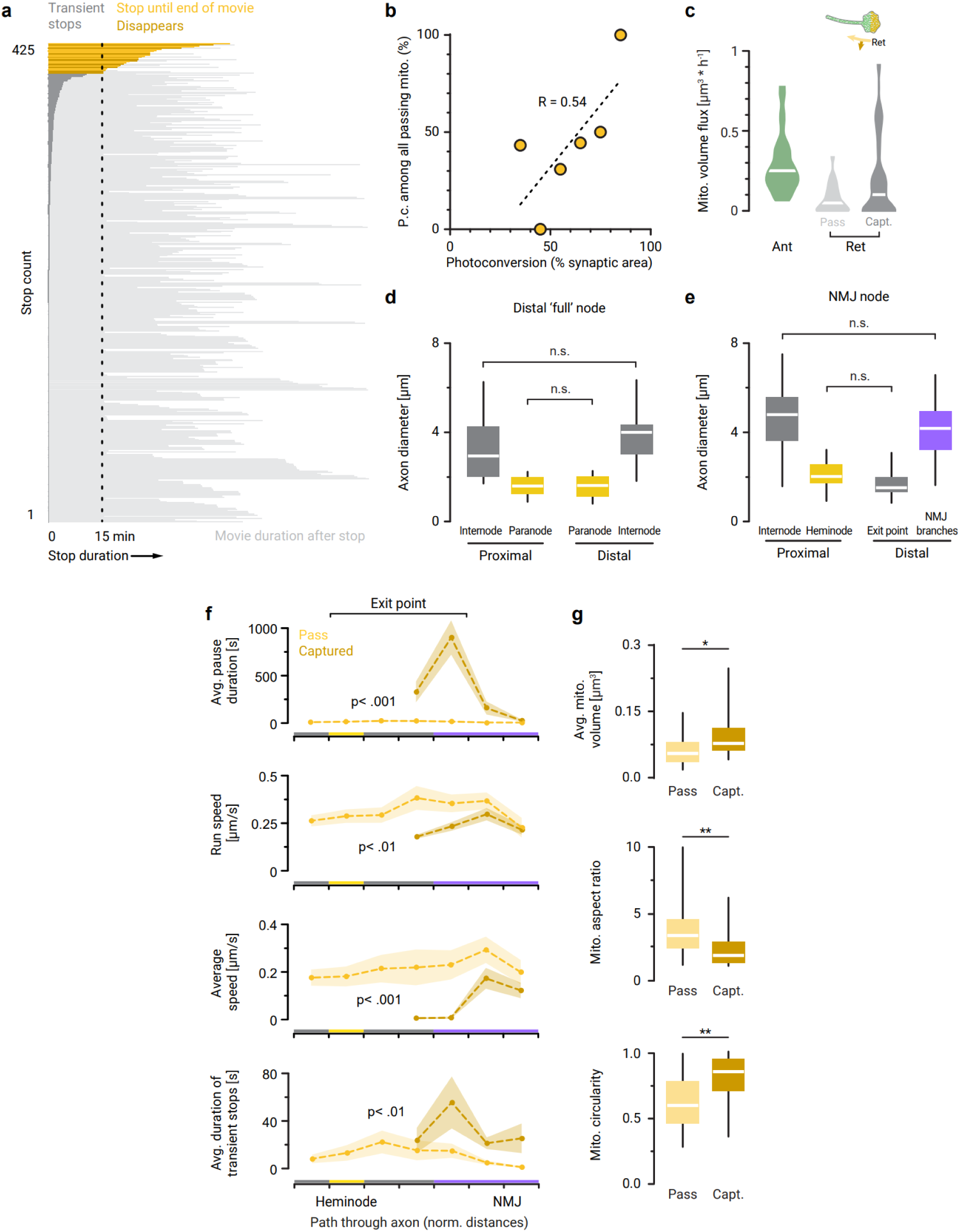
‘Captured’ mitochondria have distinguishing characteristics when compared to ‘passing’ mitochondria. **a**, Stopping behavior of retrogradely moving mitochondria in the synaptic exit point. All registered pauses (see Methods) of the photoconverted (p.c.) mitochondria that were tracked in the synaptic exit point (n = 425 stops; tracked the mitochondria in Fig. 1 **f,g** dataset) are sorted underneath each other, from longest to shortest pause. Persistent pauses (when mitochondria did not resume motility) were either observed until the end of the movie, or until the fluorescent signal slowly disappeared; see also Methods. **b**, Relationship between the fraction of synaptic area that was p.c. and the fraction of p.c. mitochondria in the overall stream of retrogradely ‘passing’ mitochondria, shown with a linear trendline (shows means per 10%-bin; n = 28 NMJs, from 19 *Thy1*-mito-Dendra mice from the Fig. 1 **f,g** dataset). **c**, Distribution of the mitochondrial volume flux data for each of the mitochondrial groups shown in Fig. 1g. **d,e**, Axonal constriction around nodes of Ranvier in α-motor axon endings (n ≥ 11 axons from triangularis sterni muscle whole mounts of ≥ 8 *Thy1*-mito-Dendra mice): **d**, calibers around the distal ‘full’ nodes, and **e**, around the synaptic exit point and its heminode (values for NMJ branches as well as for exit points show the sum of the individual branch calibers added together). **f**, Motility parameters of ‘passing’ and ‘captured’ mitochondria traveling through different axonal subcompartments (for legend, see Fig. 1f; 49 mitochondria were tracked in 21 NMJs from 16 *Thy1*-mito-Dendra mice). **g**, Shapes of ‘passing’ and ‘captured’ mitochondria (n ≥ 19 mitochondria per group, from all experiments in Fig. 1f**,g**), described by average organelle volume, aspect ratio, and circularity (see Methods). Box plots in **c,d,e,g**: median, quartile 1 - quartile 3, whiskers 95th percentiles (not shown for **c**). Graphs in **f** shown means ± s.e.m. One-way ANOVA test with multiple comparisons was used to determine significance in **d,e**, a two-way ANOVA test in **f**, and Mann Whitney U tests for **g**: ∗∗p < 0.01; ∗p < 0.05; n.s.: non-significant, p ≥ 0.05.

**Extended Data Fig. 4:**
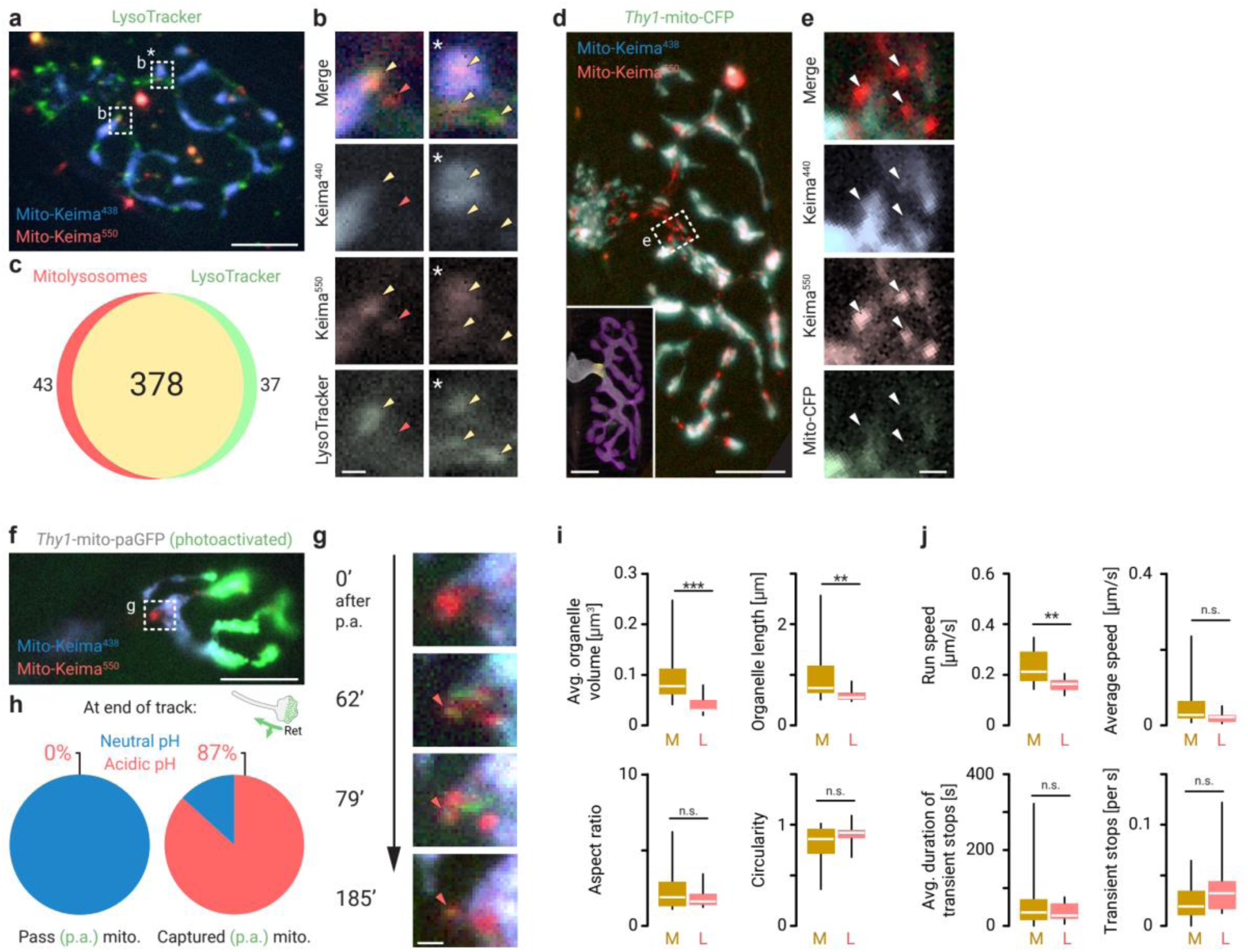
Lysosomes in motor axon terminals of acute nerve muscle explants. **a**–**c**, Colocalization of an acidophilic dye (LysoTracker, green) with mitolysosomes (mito-Keima; 438 ± 12 nm, blue and 556 ± 10 nm, red; 458 total particles analyzed from n=10 NMJs, n=3 C57BL/6 mice expressing AAV9-*hSyn*-mito-Keima). **a**, Widefield maximum projection of a neuromuscular junction, **b**, details of the two regions that are outlined in **a** by boxes, single sections of each channel showing colocalization (yellow arrows) as well as ‘no colocalization’ of a mitolysosome with LysoTracker (red arrow). **c**, Venn diagram illustrating the extent of colocalization of the two markers. **d**–**e**, Non-colocalization of a regular mito-XFP fluorophore (green) with mitolysosomes (mito-Keima; 438 ± 12 nm, blue and 556 ± 10 nm, red). **d**, Widefield maximum projection of a neuromuscular junction with a large accumulation of mitolysosomes in its exit points, *Thy1*-mito-CFP mouse labelled with AAV9-*hSyn*-mito-Keima. Inset shows post-hoc correlated staining (anti-Caspr, yellow; α-bungarotoxin, magenta; anti-βIII-tubulin, white). **e**, Widefield single optical sections of mitolysosomes in the synaptic exit point (position of image indicated by a box in **d**). The same area is shown in all three channels (blue, neutral Keima-438; red, acidic Keima-550; green, mito-CFP). Arrows point to mitolysosomes (high Keima-550 signal) that are lacking in Mito-CFP fluorescence. **f**–**h**, Colocalization of mitolysosomes and captured mitochondria over time. **f**, Mitochondria were photoactivated (p.a.) from non-fluorescent to green fluorescence in the distal half of neuromuscular junctions in acute nerve-muscle explants of *Thy1*-mito-paGFP mice expressing AAV9-*hSyn*-mito-Keima (widefield image section; blue, neutral Keima-438; red, acidic Keima-550). After p.a., the synaptic exit point (boxed area) was monitored routinely for p.a. ‘captured’ mitochondria and the proximal axon for p.a. ‘passed’ mitochondria (stacks ca. every 10–15 min to avoid photobleaching). **g**, Widefield image sections from the boxed area in **f**, taken at different time intervals indicated next to each image (min after p.a.). A ‘captured’ p.a. mitochondrion (green fluorescence) colocalized with a mitolysosome (high Keima-550/Keima-438 ratio), indicated by a red arrow in each time point. **c**, Proportion of ‘passing’ and ‘captured’ mitochondria that colocalized with mitolysosomes (red) within the duration of the experiment (n = 9 ‘passing’ and n = 15 ‘captured’ mitochondria from 10 NMJs, 5 mice). **i**,**j**, Mitochondrial (“M”) properties (analysis from the datasets in **Extended Data Fig. 3f,g** replotted, *Thy1*-mito-Dendra mice) are compared against mitolysosomal (“L”) properties (movies acquired in ≥ 4 NMJs from ≥ 3 mice per group; *Thy1*-mito-CFP mice injected with AAV9-*hSyn*-mito-Keima). **i**, Mitochondrial morphology measurements from **Extended Data Fig. 3g** are compared against measurements in mitolysosome movies (‘passing’: n = 20 mitolysosomes and ‘captured’: n = 11 mitolysosomes). **j**, Mitochondrial movement parameters from tracks in **Extended Data Fig. 3f** are compared against mitolysosome movement parameters (‘passing’: n = 22 mitolysosomes and ‘captured’: n = 8 mitolysosomes). Scale bars: **a,d,f** 10 µm; **b,e,g** 1 µm; Box plots: median, quartile 1 - quartile 3, whiskers 95th percentiles. Mann-Whitney-U test was used to determine significance in **i,j**: ∗∗∗p < 0.001; ∗∗p < 0.01; n.s.: non-significant, p ≥ 0.05.

**Extended Data Fig. 5:**
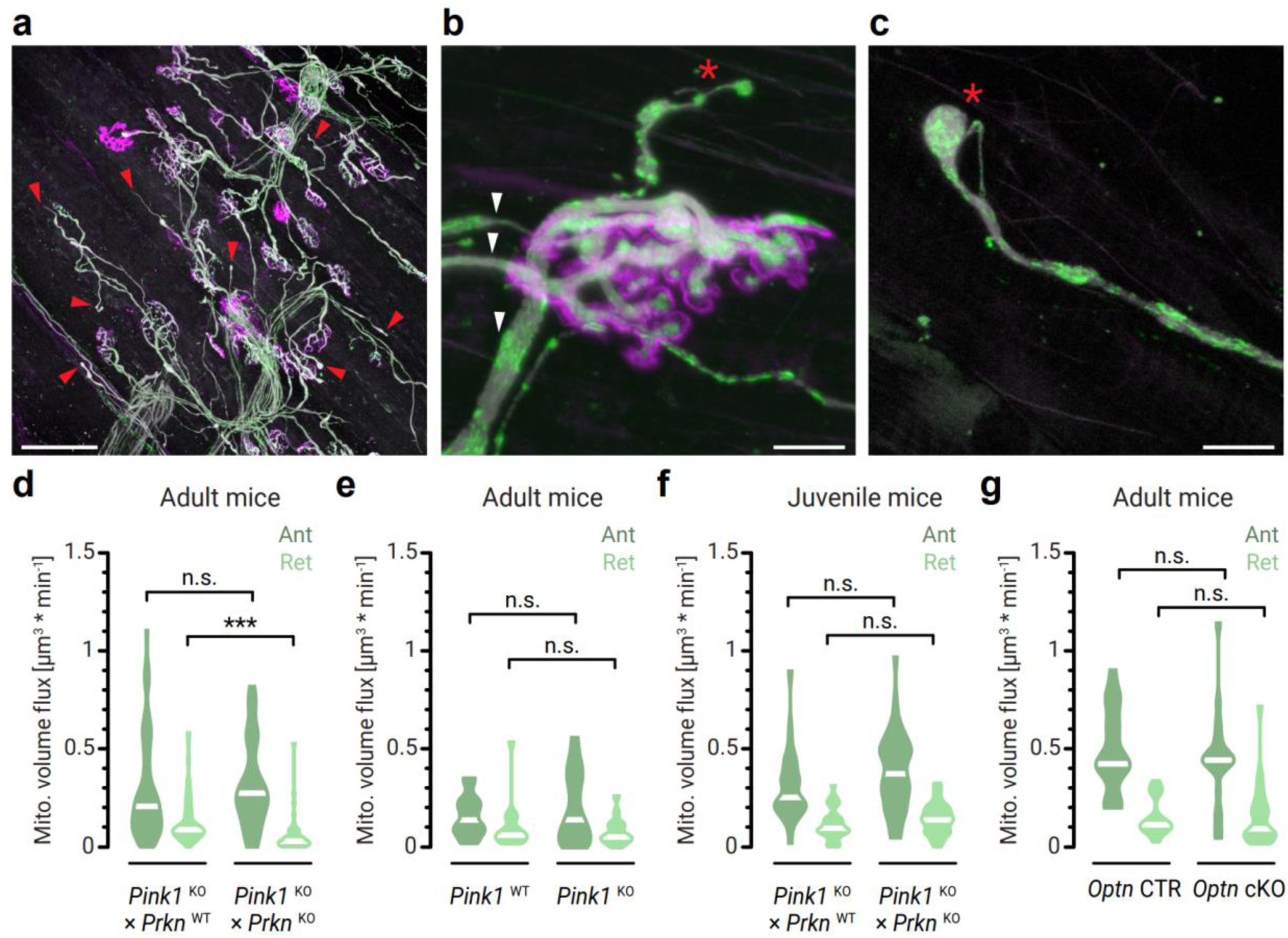
Mitophagy-proteins Pink1/parkin and optineurin in motor axon terminals of acute nerve muscle explants. **a**–**c** Neuromuscular phenotype of Pink1/Parkin-dKO mice (triangularis sterni muscles of an adult *Pink1*^ko/ko^ × *Prkn*^ko/ko^ × *Thy1*-mito-Dendra mouse; mitochondria, green; βIII-tubulin antibody, grey; AChR labelled by bungarotoxin, magenta): confocal projections show in **a** the neuromuscular endplate band, red arrows point to multiple axon sprouts (axon “stumps” that do not form a synapse), in **b** a junction that is multi-innervated (three axons indicated by arrows supply the synaptic territory), star indicates axon sprout, in **c** an axon sprout (star) with fragmented mitochondria. **d–g**, Mitochondrial volume flux measured in stem axons (intercostal nerve) of acute-nerve muscle explants in **d**, adult Pink1/Parkin-double KO (dKO) compared to single Pink1-KO (Exp: *Pink1*^ko/ko^ × *Prkn*^ko/ko^ × *Thy1*-mito-Dendra; Ctr: *Pink1*^ko/ko^ × *Prkn*^wt/wt^ × *Thy1*-mito-Dendra littermates; n ≥ 41 axons from ≥ 3 mice per group); **e**, adult single Pink1-KO compared to Pink1-WT (Exp: *Pink1*^ko/ko^ × *Thy1*-mito-Dendra; Ctr: *Pink1*^wt/wt^ × *Thy1*-mito-Dendra littermates; n = 15 axons from 2 mice per group); **f**, juvenile (3 weeks old) Pink1/Parkin-double KO (dKO) compared to single Pink1-KO (Exp: *Pink1*^ko/ko^ × *Prkn*^ko/ko^ × *Thy1*-mito-Dendra; Ctr: *Pink1*^ko/ko^ × *Prkn*^wt/wt^ × *Thy1*-mito-Dendra littermates; n ≥ 30 axons from ≥ 3 mice per group); **g**, Optn-cKO mice (right pair: cKO, ChAT-Cre^mut/wt^ × *Optn*^fl/fl^ × Thy1-mito-Dendra mice; left pair: Control, ChAT-Cre^wt/wt^ × *Optn*^fl/fl^ × *Thy1*-mito-Dendra littermate controls; n ≥ 19 axons from 3 mice per group). Scale bars in **a** 100 µm, in **a**–**c** 10 µm. Violin plots in **d,e,f,g** show data distribution with median values. Mann-Whitney-U test was used to determine significance in **d,e,f,g**: ∗∗∗p < 0.001; n.s.: non-significant, p ≥ 0.05.

## SUPPLEMENTARY MOVIE LEGENDS

**Supplementary Movie 1**

Time lapse of retrogradely moving mitochondria ‘passing’ through (blue arrow) or being ‘captured’ (cyan arrow) in the synaptic exit point (*Thy1*-mito-Dendra labels all mitochondria in green; photoconverted mitochondria in red). Movie from the neuromuscular junction in **Fig. 1d,e**. Latter half of the movie shows the overlying manual trackings.

**Supplementary Movie 2**

Photoconverted mitochondria (red) moving retrogradely through the last ‘full’ node of Ranvier (yellow, initial frame) in of a motor neuron terminal (*Thy1*-mito-Dendra mouse line).

**Supplementary Movie 3**

Photoconverted mitochondria (red) moving anterogradely through the last ‘full’ node of Ranvier (yellow, initial frame) in of a motor neuron terminal (*Thy1*-mito-Dendra mouse line).

**Supplementary Movie 4**

Photoconverted mitochondria (red) moving retrogradely through node of Ranvier (yellow, initial frame) in the stem axon of the intercostal nerve (*Thy1*-mito-Dendra mouse line).

**Supplementary Movie 5**

Murine neuromuscular junction labelled with AAV9-*hSyn*-mito-Keima; Mitolysosomes appear high in acidic signal (red), mitochondria appear high in neutral Keima-signal (blue). Manual trackings in the latter half of the movie follow bidirectionally moving mitolysosomes.

## ONLINE METHODS

### Animal models

We performed live imaging of neuronal mitochondria in transgenic mouse lines that express mitochondria-tagged fluorescent proteins under the neuronal *Thy1*-promoter (termed *Thy1*-mito-XFP lines; XFP stands for either CFP, RFP, or Dendra2) (Misgeld et al., 2007; Breckwoldt et al., 2014). Subpopulations of mitochondria were photo-labeled with a localized UV-light pulse, using newly generated *Thy1*-mito-Dendra and *Thy1*-mito-paGFP mouse lines (see next section, cf. Magrané et al., 2014). Since paGFP is non-fluorescent in its native state (Patterson & Lippincott-Schwartz, 2002), we crossed *Thy1*-mito-paGFP to *Thy1*-mito-RFP mice (Breckwoldt et al., 2014) to visualize neuronal mitochondria in the non-photoactivated area. Transgenic mice with expression of a fluorescent protein in neuronal cytoplasm (*Thy1*-YFP-16 mice) (Feng et al., 2000) were used to colocalize LysoTracker to the axoplasm. ChAT-Cre crossed to ROSA-mito-GFP mice were used to visualize mitochondria in motor neurons for TMRE incubation experiments (Cre enzyme expression in cholinergic neurons driving mito-GFP expression; Agarwal et al., 2017; Rossi et al., 2011). Experimental models were crossed to *Thy1-*mito-Grx1-roGFP2 mice (Breckwoldt et al., 2014) to measure mitochondrial reactive oxygen species (ROS) levels. C57BL/6 mice (The Jackson Laboratory; Stock #000664) or transgenic mice expressing a fluorophore under the *Thy1*-promoter were used for AAV-injections (indicated in figure legends).

To elucidate the molecular mechanism of mitochondrial quality control in axon terminals, we used the following knock-out (KO) mice and controls: ChAT-Cre^mut/wt^ × *Atg7*^flox/flox^ × *Thy1*-mito-Dendra vs. ChAT-Cre^wt/wt^ × *Atg7*^flox/flox^ × *Thy1*-mito-Dendra and ChAT-Cre^mut/wt^ ^or^ ^wt/wt^ × *Atg7*^wt/wt^ × *Thy1*-mito-Dendra controls (“*Atg7*-cKO”, Komatsu et al., 2005); *Pink1*^-/-^ × *Thy1*-mito-Dendra vs. *Pink1*^+/+^ × *Thy1*-mito-Dendra littermate controls (“*Pink1*-KO”, Glasl et al., 2012); *Pink1*^-/-^ × *Prkn*^-/-^ × *Thy1*-mito-Dendra vs. *Pink1*^-/-^ × *Prkn*^+/+^ × *Thy1*-mito-Dendra littermate controls (“*Pink1-Parkin*-DKO”, Itier et al., 2003); ChAT-Cre^mut/wt^ × *Optn*^flox/flox^ × *Thy1*-mito-Dendra vs. ChAT-Cre^wt/wt^ × *Optn*^flox/flox^ × *Thy1*-mito-Dendra littermate controls (“*Optn*-cKO”, Munitic et al., 2013). We further employed two animal models of Amyotrophic lateral sclerosis (ALS). The following experimental and control groups were used: SOD1^G93A^ ^+/-^ (“mut”) × *Thy1*-mito-Dendra vs. SOD1^G93A-/-^ (“wt”) × *Thy1*-mito-Dendra littermate controls (Gurney et al., 1994); *Thy1*-(GA)149-CFP ^+/-^ (“mut”) × *Thy1*-mito-Dendra vs. *Thy1*-(GA)149-CFP ^-/-^ (“wt”) × *Thy1*-mito-Dendra gender-matched littermate controls (Schludi et al, 2017). Each SOD1^G93A^ ^+/-^ breeder was monitored and scored weekly for the development of motor symptoms.

Tail or ear biopsies were genotyped with standard PCR protocols using the primers in **Extended Data Table 1**. For all experiments, mice of both sexes were used between the ages of 6 weeks to 24 months, with the exception of the following datasets, where we restricted data acquisition to ages before phenotype onset: *Pink1*-KO and *Pink1-Parkin*-DKO datasets in **Fig. 5a–d** and **Extended Data Fig. 5f** (postnatal days 21–27), SOD1^G93A^ (postnatal days 14–21), and *Thy1*-(GA)149-CFP breedings (5–6 weeks).

### Generation of *Thy1*-mito-paGFP and *Thy1*-mito-Dendra transgenic lines

We used the *Thy1*-promoter to drive the expression of mito-Dendra and mito-paGFP selectively in neurons as previously described (Marinkovic, Godinho and Misgeld, 2015a,b,c; Misgeld et al., 2007). In brief, to create mito-Dendra and mito-paGFP fusion genes, an N-terminal in-frame fusion between the coding sequence of the fluorescent protein (Dendra2 or paGFP) (Gurskaya et al., 2006; Patterson and Lippincott-Schwartz, 2002) and the mitochondrial targeting sequence from subunit VIII of the human cytochrome c oxidase gene (Clontech) was generated. After cloning the fusion gene into the *Thy1* vector, standard pronuclear injection was performed to produce several independent founder lines, which were screened for expression patterns, levels and photoswitching properties. Two bright and broadly expressing lines were retained and used for further experiments (**Extended Data Table 2**, **Extended Data Fig. 1**). Homozygotes of all lines bred normally and showed no neurological abnormalities.

### Widefield imaging of mitochondrial dynamics in axons from acute nerve-muscle explants

Triangularis sterni muscle explants were dissected from mice using a previously described protocol with modifications (Kerschensteiner et al., 2008; Brill et al., 2013). Mice were lethally anaesthetized with isoflurane (Abbott) and explants were prepared under a dissection microscope (Olympus SZ51). We dissected *Thy1*-mito-Dendra and *Thy1*-mito-paGFP mice in a dark room under red light to avoid premature photoswitching. Briefly, the thorax was exposed by removing skin and pectoral muscles and released from the surrounding tissue by severing the ribs close to the spine in < 1 min. The explanted thorax, which contains the triangularis sterni muscle and its innervating intercostal nerves, was transferred to a dish with oxygenated (bubbled with 95% O_2_ / 5% CO_2_) and cooled Ringer’s solution on ice (in mM: 125 NaCl, 2.5 KCl, 1.25 NaH_2_PO_4_, 26 NaHCO_3_, 2 CaCl_2_, 1 MgCl_2_, and 20 glucose), and any remaining inner organs were removed. The explant was pinned on a Sylgard-coated 3.5 cm dish with insect pins (Fine Science Tools, 26002-20), the ventral side with the triangularis sterni muscle facing up. During imaging, the explant was continually perfused with warmed, oxygenated Ringer’s solution at 2–3 mL / min and kept at temperatures between 33–36 °C in a heating ring (Warner instruments). Organelle transport and dynamics were recorded on µManager-controlled epifluorescence microscopes (Olympus BX51WI; www.micro-manager.org), equipped with ×4/0.5 N.A. air-, ×20/0.5 N.A. and ×100/1.0 N.A. water-immersion dipping cone objectives, mercury arc lamp (Olympus) or LED light source (pE-4000 CoolLED), 405 nm LED (Thor Labs, #M405L1) or laser (Coherent), automated filter wheel (Sutter), CCD (Retiga EXi, Qimaging) or CMOS camera (Andor Sona, low-noise mode) and motorized stage (MMTP, Scientifica; Ludl, MAC 6000). Neutral density and infrared-blocking filters were used in the light path to prevent phototoxicity. The following multicolor filter sets were used: Triple-FITC/Cy3/Cy5 (Chroma #86016) or LED-DA/FI/TR/Cy5-B-000 (Semrock) for mito-Dendra; Triple-ECFP/EYFP/mCherry (Chroma #89006) or LED-CFP/YFP/mCherry-A-000 (Semrock) for mito-RFP and mito-paGFP; and GFP/Keima(pH)-3X2M-A-000 (Semrock). For axonal transport measurements in intercostal nerves (stem axons), movies of 180–300 images were acquired at 1 Hz and exposure time of 300 ms. At least three movies were recorded per experiment. For synaptic transport measurements, movies were acquired for 35 min–2.5 h (0.5–0.33 Hz) at the “entry points” of terminal axon branches into superficial neuromuscular junctions (NMJs) innervating the triangularis sterni muscle. No more than two NMJs were imaged per explant to keep the imaging time window within the limit of physiological transport rates (∼3 h). Low-magnification stacks (×20 and ×4) were taken to enable re-localization of recorded synapses after sample fixation and immunohistochemistry.

### Photoswitching

Photoactivation / -conversion was performed with 405 nm illumination using a ×100/1.0 N.A. water-immersion dipping cone objective and restrained to the area of interest with an aperture stop (Olympus; resulting spot on the sample ∼40 µm in diameter). For pulse–chase experiments, the smallest dose of 405 nm irradiance that yielded adequate contrast was used to minimize potential phototoxicity effects (Dendra ∼870 µW*s, pa-GFP ∼8.7 mW*s; ca. 30-fold increase in fluorescence intensity, see **Extended Data Fig. 1c,g**, resulting in fluorescence intensities of 405-nm-illuminated mitochondria more than 4 standard deviations away from the mean of non-illuminated mitochondrial populations, **Extended Data Fig. 1d,h**).

### Mito-photo-DNP incubation and TMRE staining

For depolarization experiments in *Thy1*-mito-Dendra mice, acute nerve-muscle explants were incubated with mito-photo-DNP (Chalmers et al., 2012; Focus Biomolecules, 10-1580) in a dark room under red light as a bath application for 30 min (using the perfusion system and widefield live imaging setup; diluted at 20 µM in prewarmed oxygenated Ringer’s solution; Glancy et al., 2015). In the experimental group, mito-photo-DNP was photoactivated immediately after washout (7 min) using a ∼870 µW*s UV dose, while in the ‘no depolarization’ control group, the UV dose was applied before start of the mito-photo-DNP incubation, resulting in photoconversion of mito-Dendra for both groups, but photoactivation of mito-photo-DNP only in the experimental group. To measure mitochondrial membrane potential under photo-mito-DNP depolarization, acute explants were dissected from ChAT-Cre × ROSA-mito-GFP mice for live imaging and perfused with 30 nM TMRE at 30–33 °C (non-quenching condition; diluted in oxy-Ringer’s with 0.1% DMSO; Thermofisher, T669). Stacks of junctions that were located in successfully TMRE-labelled muscle tissue were used for experiments: neuromuscular junctions after photoactivation of mito-photo-DNP (experimental condition) and with only UV-illumination (control) were acquired.

### Generation of recombinant DNA

mKeima-Red-Mito-7 was a gift from Michael Davidson (Addgene plasmid # 56018) and pAAV-hSyn-DIO-HA-hM3D(Gq)-IRES-mCitrine was a gift from Bryan Roth (Addgene plasmid # 50454). To generate the *hSyn*-mito-Keima construct for AAV9-production, mito-Keima was digested from the mKeima-Red-Mito-7 plasmid using *NheI* and *NotI* and ligated into a self-complementary AAV backbone plasmid. The human Synapsin promoter was amplified by PCR from the pAAV-hSyn-DIO-HA-hM3D(Gq)-IRES-mCitrine plasmid and then cloned before the mKeima-Red-Mito-7 using *MluI* and *NheI*.

### Generation of AAV9

For AAV9-hSyn-mito-Keima production, HEK293-T cells were grown in 10-tray Cell Factories (Thermo Fisher Scientific) using Dulbecco’s modified essential medium (DMEM) containing 10% fetal bovine serum (Gibco) and 1% penicillin/streptomycin for 24 h before transfection. The transgene plasmid (420 µg of ‘pAAV-*hSyn*-mito-Keima’) and the helper plasmid (1.5 mg of pDP9rs, kindly provided by Roger Hajjar, Icahn School of Medicine at Mount Sinai, New York) were transfected into the HEK293-T cells using polyethylenimine (Polysciences). 72 h later, the cells were harvested, lysed, and benzonase-treated. The virus particles were purified by ultracentrifugation through an iodixanol density gradient (Optiprep, Progen). Iodixanol was replaced by Ringer lactate buffer (Braun) using Vivaspin 20 columns (Sartorius). The latter also aimed at concentrating the virus by decreasing the volume of the solution. Viral titers were determined by real-time PCR using SYBR Green Master Mix (Roche) to be in the range of 1 × 10^13^–2 × 10^14^ genome copies per milliliter.

### AAV injection into the central nervous system of neonatal mice

AAV were injected into mice as described previously (Wang et al., 2021). Briefly, pups were anaesthetized with ∼32% isoflurane between postnatal days 2–3 and injected with 2 µL of AAV-particle suspension into the brain ventricle, under ultrasound guidance (Vevo1100 Imaging System and Microscan MS550D transducer, Visualsonics) and with a glass pipette attached to a nanoliter injector (30 nL/s; Micro4TM MicroSyringe Pump Controller and Nanoliter 2000, World Precision Instruments; pipette: 3.5’’ Drummonds #3-000-203-G/X). Afterwards, mice recovered on a heating mat (37 °C) and were returned to the mother. After 6 weeks of age, injected mice were sacrificed for live explant imaging as described above.

### Confocal live imaging: LysoTracker, TMRM, mitoROS

To determine LysoTracker distribution in motor axon terminals, acute triangularis sterni nerve-muscle explants of *Thy1*-YFP-16 mice were dissected as described above and transferred to an upright confocal setup equipped with ×100/1.0 N.A. and ×20/.5 N.A. water-immersion objectives, a heated stage, and perfusion system (Olympus FV1000; 33–36°C; 95% O_2_, 5% CO_2_ bubbled Ringer’s solution). The explants were incubated with LysoTracker Red DND-99 for 3 min (1:5000, Invitrogen, L7528) and Alexa-647-conjugated α-bungarotoxin for 15 min (1:50, Invitrogen, B-35450, 50µg/ml) added to perfusion media (Song et al., 2008). After washing, confocal image stacks of superficial axon terminals were acquired at Nyquist frequency, and positions of the terminals were mapped (×20 objective) to enable re-localization after sample fixation. Mitochondrial membrane potentials and ROS levels from explants of Atg7-cKO mice and controls were acquired using a similar confocal live setup. For membrane potential measurements, acute explants were dissected and perfused with 30 nM TMRM at 30–33 °C (non-quenching condition; diluted in oxy-Ringer’s with 0.1% DMSO; Thermofisher, T668), and stacks of superficially located junctions were acquired.

### Fixed sample preparation and immunohistochemistry

For immunohistochemistry, thoraces were fixed in 4% paraformaldehyde (PFA; Sigma #P6148 or Electron Microscopy Sciences #15710), or 2% PFA for anti-ubiquitin stainings, in 0.1 M phosphate buffer (PB; Na_2_HPO_4_/NaH_2_PO_4_, pH 7.35–7.40), 1–2 h on ice. Triangularis sterni muscles were dissected as whole mounts to leave the microanatomy of NMJs, distal axon branches and innervated myocytes intact (Brill et al., 2013; Kerschensteiner et al., 2008). For some antibody stainings (anti-ubiquitin, anti-Dendra, anti-GFP), muscles were additionally permeabilized for 1 h at 37 °C with 5% CHAPS (300 rpm; Roth #1479.3). For antibodies raised in mouse, muscles were additionally blocked with mouse Fab-fragments (1:100 in PB), then blocking solution (see below), with each incubated for 1 h on a shaker (60 rpm) at room temperature. Brains and spinal cords were collected from mice following lethal anesthesia with isoflurane and transcardial perfusion with ice-cold4 % PFA in 0.01 M phosphate-buffered saline (PBS; pH 7.35). After immersion in fixative overnight (4°C), samples were mounted in 2% agarose/PBS and cut on a vibratome (Leica) into 60 µm sections.

Fixed samples were incubated overnight at 4 °C 60 rpm in the respective primary antibodies diluted in blocking solution (Muscle: 10% goat serum Millipore #S26, 1% BSA Sigma Aldrich A9647, 0.5% Triton X-100 Sigma-Aldrich T9284, in 0.1 M PB; Brain and spinal cord: 10% goat serum Sigma Aldrich G9023, 1% BSA Sigma Aldrich A9647, 0.5% Triton X-100 Sigma-Aldrich T9284, 0.05% NaN_3_, Riedel de Haen 13412, in 0.01 M PBS). The following primary antibodies were used in this study: anti-βIII-tubulin conjugated to Alexa-488 (mouse IgG2a, 1:200; Biolegend, #801203), to Alexa-555 (mouse IgG2a, 1:100; BD Biosciences #560339), to Alexa-594 (mouse IgG2a, 1:200; Biolegend #657408), or to Alexa-647 (mouse IgG2a, 1:200; Biolegend #657406); anti-Caspr (rabbit polyclonal, 1:500, Abcam, ab34151); anti-Dendra2 (rabbit polyclonal, 1:500, Evrogen, AB821); anti-GFP (rabbit polyclonal, 1:1000, Abcam, ab13970); anti-ubiquitin clone Ubi-1 (mouse IgG1, 1:10-1:20, Millipore, MAB1510). To label postsynaptic nicotinic acetylcholine receptors, Alexa-594-, Alexa-647- or biotin-conjugated α-bungarotoxin (1:20, Invitrogen, B-13423, B-35450, B1196; 50µg/mL in PBS) was added to the primary antibody mixture. Brain and spinal cord sections were stained with Neurotrace 640/660 (Invitrogen N21483), diluted 1:500 in the primary antibody solution. Samples were washed in 0.1 M PB (muscles) or 0.01 M PBS (vibratome sections), incubated for 2 h on a shaker (60 rpm) at room temperature with streptavidin coupled to Alexa-405 (1:200, Invitrogen, S32351; 1mg/mL in PBS) and/or suitable secondary antibodies coupled to Alexa-Fluor dyes (raised in goat; Thermofisher and Jackson Immunoresearch), then washed again. Samples were mounted on glass slides in Vectashield (Vector Laboratories) or Fluoromount G (Thermofisher, #00-4958-02). Muscles were flattened by placing them under pressure between glass slide and cover slip.

### Confocal and Airy scan microscopy

Image stacks of fixed samples were recorded on a confocal microscope equipped with ×10/0.40 N.A. air-, ×20/0.8 N.A., ×40/1.35 and ×60/1.42 N.A. oil-immersion (Olympus FV1000, Olympus FV3000), or ×10/0.45 N.A. air-, ×20/0.8 N.A. air-, and ×63/1.4 N.A. oil-immersion objectives (Zeiss LSM980). NMJs that were live imaged before fixation were re-localized in the fixed sample based on anatomical fiducials (innervation and axon branching patterns, position within the muscle). All samples were acquired using Nyquist sampling, sequential scanning, and automated pinhole diameter. Samples imaged by Airy Scan microscopy (LSM 800; LSM 980 with Airy Scan 2; Zeiss) were mounted using ultra fine coverslips (Hecht Assistent, #41014521). Laser power and detector voltage were adjusted to neither under-nor oversaturate the signal; for quantitative datasets, settings between experiments and controls were kept constant for comparability. Airy Scan stacks were processed with Zeiss ZEN Blue software using appropriate Wiener filter parameters, with settings between experiments and controls kept constant.

### Serial and correlative electron microscopy

The *M. Triangularis sterni* explant from an 11-week-old *Thy1*-YFP-16 mouse was prepared for confocal live imaging, stained with LysoTracker and α-bungarotoxin-647 and mapped as described above. Immediately after confocal stack acquisition from superficial NMJs, explants were transferred to cooled EM fixative (4% PFA, Electron Microscopy Sciences; 2.5% glutaraldehyde, Electron Microscopy Sciences; in 0.1 M sodium cacodylate; 0.45 µm filtered) for 24 h on 4 °C. The thorax region containing the imaging site was punched (2 mm diameter) and transferred to a glass imaging slide. NMJs were re-localized and outlined by two asymmetrically shaped boxes using two-photon laser burns (near-infrared branding “NIRB”; Bishop et al., 2011) on an Olympus FVMPE-RS microscope equipped with a ×25/1.05 N.A. water-immersion multiphoton objective and GASP detectors (NIRB settings: 100% laser power, 920 nm, 30–60 s). After post-fixation (48 h at 4 °C), NIRB marks were re-localized and the triangularis sterni muscle layer dissected before embedding. A standard rOTO protocol was applied for *en bloc* staining (Kislinger et al., 2023). After post-fixation with reduced osmium (2% osmium tetroxide, 1.5% potassium ferrocyanide in 0.1 M sodium cacodylate buffer pH 7.4), TCH treatment (1 % aqueous thiocarbohydrazide; Sigma, 223220-5G) another osmium step (2% aqueous osmium) followed. The tissue was incubated overnight in 1% uranyl acetate at 4°C, contrasted in 0.1% lead aspartate (silver nitrate, Alfa Aesar, A16345; L-aspartate, Sigma, A9256-100g), dehydrated, infiltrated with epon resin (Serva) and embedded in a support tissue scaffold (mouse cortex) to avoid wrinkle formation. The embedded tissue was trimmed by 200 µm to expose a rectangular block face using a TRIM90 knife (Diatome, Trim 90, DTB90) on a PowerTome ultramicrotome (RMC). Consecutive sections were taken on an automated tape collecting ultramicrotome (ATUM) with a diamond ultra-knife (Diatome) at 50 nm thickness and collected on carbon-coated Kapton tape (kindly provided by Richard Schalek, Jeff Lichtman, Harvard; Schalek et al., 2012). The tape was plasma-treated (custom-built vacuum chamber based on the glow discharger easiGlow, PELCO) right before section collection (Kasthuri et al., 2015; Kislinger et al., 2023). The appearance of NIRB marks was monitored directly during sectioning and by regular sampling and light microscopic inspection of sections on tape after methylene blue staining (Sigma, M9140). Kapton strips bearing the selected sections were assembled onto double-sided adhesive carbon tape (Science Services, P77819-25), mounted onto a 4-inch silicon wafer (Science Services, SC4CZp-525-01) and grounded with adhesive carbon tape strips (Science Services, 77816-25). Section images were acquired on a Crossbeam Gemini 340 SEM (Zeiss) in backscatter mode at 8 keV (high gain) at approximately 7 mm WD and 60 µm aperture. Wafers were screened for NIRB marks and selected for acquisition. In ATLAS5 Array Tomography (Fibics, Ottawa, Canada) the entire wafer was imaged at 6000 nm/pixel followed by mapping and medium resolution (200 nm/pixel) imaging of individual tissue sections. The region of interest was automatically acquired at 3 nm/pixel. The images were aligned semiautomatically using *Fiji* TrakEM2 Plugin (Cardona et al., 2012) and annotated in IMOD for rendering (https://bio3d.colorado.edu/imod/; Kremer et al., 1996). A similar procedure (excluding the confocal live imaging and NIRB correlation routine) was performed for the dataset in **Fig. 4b,d** (Atg7-cKO and control mouse).

### Data analysis

Image analysis was performed with the open-source ImageJ/Fiji (http://fiji.sc) (Schindelin et al., 2012). To account for sample drift, movies from live explants were aligned to a region of interest prior to analysis (‘Template matching’ plugin, Tseng et al., 2011). Fluorescence intensity values were gained by measuring the mean pixel value in a region of interest (ROI), followed by subtraction of the mean background pixel value in the corresponding frame or stack.

### Fluorescence intensity and UV-dose–response curves

To gain fluorescence intensity measurements of photoswitched and non-photoswitched forms of paGFP and Dendra, individual mitochondria were isolated in neuromuscular junctions (NMJs) by manually drawn ROIs. At least three measurements were taken per mitochondrion and their average fluorescence intensity calculated in Fiji for each color channel. For dose-response plots, NMJs were illuminated with incremental steps of UV light and mean values from each step were plotted against the corresponding cumulative duration of UV illumination. The measurements were normalized to the initial values before UV illumination. To account for sample drift, the ROIs were repositioned manually to the selected mitochondria. In addition, the ratios of photoswitched (p.s.)/non-photoswitched (non-p.s.) fluorescence intensities were plotted in a histogram at the dosage that was deemed useful for subsequent live imaging experiments (Dendra: ∼870 µW*s; paGFP: ∼8.7 mW*s UV light). There, the photo-labeled were differentiated from non-labeled mitochondria based on their p.s./non-p.s. ratios: a mitochondrion was considered to be photo-labeled if the ratio exceeded the mean prior to UV illumination by 4 standard deviations (**Extended Data Fig. 1**).

### Mitochondrial transport and shape

In Airy Scan image stacks, mitochondrial volume was calculated by outlining a mitochondrion’s area in each z-section and multiplying the area sum with section thickness. A mitochondrion’s area was defined as all cohesive pixels with intensity surpassing ½ of the peak intensity in the corresponding line scans (‘Plot Profile’ *Fiji* function, example in **Extended Data Fig. 2g,h**), multiplied by the resolution in xy. In correlated widefield stacks, mitochondrial length (*l*) and width (*w*) were measured, as defined by the distance between the ½ maxima of fluorescence intensity in a line scan (**Extended Data Fig. 2c,d**). Mitochondrial volume was then calculated by assuming an ellipsoidal shape: 1/6 *π* ∗ *l* ∗ *w*^2^.

In time-lapse movies, fluorescent organelles that passed through an axonal cross-section in either anterograde or retrograde direction were manually annotated (Misgeld et al., 2007; Marinkovic et al., 2012). A trained observer measured mitochondrial length (*l*) and width (*w*) with the ‘line’ tool in *Fiji* to calculate their volume according to the above formula. For area and perimeter, mitochondria were outlined with the ‘freehand selection’ tool. In axon terminals, each mitochondrion was measured in three different time-lapse frames if possible, and the average used for further analysis. Mitochondrial volume flux rates were calculated as the sum of transported mitochondrial volume divided by observation time. Mitochondrial aspect ratio was given as *l* divided by *w*. Circularity is calculated as 4 *π* ∗ area / perimeter^2^. Both shape factors represent a perfectly circular shape by the value of 1.

### Analysis of organelle motility

The ‘Manual Tracking’ *Fiji* plugin was used to track the movement of mitochondria labelled by photoconverted mito-Dendra and the movement of mitolysosomes labelled by mito-Keima. A table was extracted with coordinates (x,y) for each time frame where an organelle was registered. After computing the Euclidian distance between consecutive positions, each track was divided into sections of motility vs. pausing behavior. Stops (or pauses) were registered where a previously motile organelle did not move or was displaced by less than 2 pixel per frame (∼0.09 µm/s). If an organelle resumed motility after a stop, this was registered as a ‘transient’ stop. Transient mitochondrial stops did not last for longer than 15 min (**Extended Data Fig. 3a**); instead, mitochondria that stopped for longer than 15 min stayed in their position until the end of the movie or until they slowly disappeared (**Extended Data Fig. 3a**). These stops were marked as ‘persistent’ stops. Next, the following motility parameters were calculated for each organelle: average speed (total distance travelled divided by total observation time), run speed (the same, but after exclusion of time spent stopping), average pause duration (total time spent pausing divided by pause count), average duration of transient stops (the same, but excluding all stops that were at the start or end of the track), and average pause rate (total number of stops divided by observation time).

Next, each mitochondrial track was mapped to the anatomy of its axon terminal. The axon outline was annotated in each movie (based on a correlated staining of βIII-tubulin) and then divided into synaptic region (α-bungarotoxin), paranodal areas (anti-caspr), internodes and unmyelinated axon (only βIII-tubulin) along its length. These distinct compartments were sectioned further to account for their varied dimensions (from distal to proximal: 10 bins between the base and the tips of the synaptic branches, 10 bins along the internodal length, 2 bins in between heminode and synapse). Each mitochondrial position in time was assigned to a bin based on proximity, therefore allowing us to plot mitochondrial tracks and motility parameters relative to their position along axonal landmarks. The likelihood whether a mitochondrial track would end in the bin due to a permanent stop (defined as a mitochondrion not moving away in the remaining observation time; only mitochondria were included that stopped at least 15 min before the end of the movie) was calculated after annotating whether a mitochondrion passed the bin (Value 0) or whether the mitochondrion permanently stopped there (Value 1), and the mean plotted as a probability over different compartments (**Fig. 1f**).

### Mitochondrial capture analysis

Mitochondrial ‘captures’ were defined as events, where retrogradely moving mitochondria stopped persistently (> 15 min) in the synaptic exit point (the region between the proximal 1/10 of the presynapse and the heminode, corroborated by post-hoc staining where possible). In contrast, ‘passing’ mitochondria traversed this region and entered the axonal internode. Pie charts show the relative proportion of photoconverted mitochondria that were either captured or passing mitochondria according to this definition (excluding mitochondria which passed within the last 15 min of the movie). “U-turns” were not included in this calculation, i.e. events where retrogradely moving mitochondria changed their direction again to return back to a more distal location within the synapse; such events were rare (∼5% of retrogradely moving mitochondria) and were not particularly associated with the synaptic exit point.

For mitochondrial volume flux rates, all ‘passing’ and ‘anterogradely’ moving mitochondria were annotated (both p.c. and non-p.c. organelles were observed using the ‘Merge’ color mode in *Fiji*) and analyzed as detailed above. ‘Captured’ mitochondria were also annotated, but the last 15 min of the movie were excluded from their volume flux rate calculation. The flux of ‘non-photoconverted’ captured mitochondria was estimated per junction from the photoconverted flux rate by normalizing for the relative proportion of the synaptic area that was photoconverted (**Extended Data Fig. 3b**). Synaptic areas were measured by thresholding the mitochondrial signal (‘Otsu’ algorithm, or ‘Li’ for the dimmer fluorescence in juvenile animals) in maximum projected, deconvolved (‘Subtract background’ *Fiji* Plugin) widefield image stacks of the axon terminals before and after photoconversion.

### Measurement of axon caliber

Using anti-βIII-tubulin stainings to infer axon outlines, axon diameters were manually measured in confocal maximum projections using the ‘line tool’ in *Fiji*. In Atg7-cKO and control datasets (**Fig. 4l**), the axon calibers were measured at the widest position in each internode.

### Measurement of mitochondrial reactive oxygen species (ROS)

Mitochondrial ROS levels in neuromuscular junctions were analyzed from confocal stacks that were acquired in acute nerve-muscle explants from mice expressing the ratiometric marker *Thy1-*mito-Grx1-roGFP2 (Breckwoldt et al., 2014). In brief, synaptic mitochondria were manually outlined in *Fiji* with ROIs, and the 405/488 ratio was calculated using the mean pixel values extracted from the 405nm- and 488nm-excitation channels (after background pixel value subtraction).

### Mito-Keima ratiometric image analysis and LysoTracker distribution

To measure mitolysosome distribution in mito-Keima expressing neurons, a combined segmentation mask was generated from the 438- and 556-nm channels after deconvolution (‘Subtract background’ Fiji Plugin; to remove unfocused scatter light), manual thresholding and binarization. The mask was used to retain the pixels containing mito-Keima signal in the stack (after background subtraction). The resulting image stacks were divided (556/438-nm) and flattened by averaging the pixel values inside the mask across the z-dimension.

To measure the distribution of LysoTracker staining in *Thy1*-YFP-16 mice, the channel with cytoplasmic YFP signal was thresholded manually and binarized to produce a segmentation mask. This mask was used on the LysoTracker channel to retain signal with intra-axonal localization and remove the staining of the surrounding tissue from the analysis. After background subtraction, the masked image stack was flattened by averaging the retained pixel values across the z-dimension.

The axon terminals from both datasets were sectioned into subcompartments by help of a correlated axon compartment staining (described above), and pixel intensity profiles were collected from each compartment by longitudinal line-scans (‘Plot Profile’ function in Fiji; the width of each annotation line was individually adjusted to fully cover the respective axon branch). The results were binned according to their relative position within the subcompartments and averaged in each bin (for LysoTracker plots, the results were additionally normalized to the overall mean value of the internodal compartment).

### Ubiquitin content in motor axon terminals

In axon terminals stained with antibodies against ubiquitin, mean pixel intensity values were extracted from confocal single optical sections where ROIs had been manually drawn in Fiji around proximal (incl. synaptic exit points) and distal parts (towards the tips) of the NMJ branches. The ROIs were outlined using the mito-Dendra channel (without viewing the ubiquitin staining) to avoid bias. For background measurements, ROIs were drawn in close proximity around the corresponding regions and their mean intensity values subtracted from the calculation.

### Image Representation

Different channels of an image series were combined in pseudo-color using Adobe Photoshop (‘screen’ function). Maximum intensity projections were generated using Fiji, and where needed, stitched in Adobe Photoshop. For figure presentation, images show maximum intensity projections (Fiji) of confocal or widefield image stacks; otherwise, figure legends indicate if single optical sections are shown. For presentation purposes, widefield stacks were deconvolved (*Subtract background* Plugin in Fiji) to remove unfocused scatter light prior to maximum projection. In non-quantitative panels, gamma settings was adjusted non-linearly to enhance visibility of low-intensity objects; gamma settings remained linear to present images of ratiometric fluorophore quantification (mito-Keima, ubiquitin staining). Images were placed on dark monochrome backgrounds where appropriate, with the boundary visible due to a non-clipped noise background. For presentation only, movies were denoised using the ‘Candle’ algorithm in MATLAB and bleaching-corrected in Fiji using an exponential fit. Renderings of segmentations from electron microscopy stacks were made in AMIRA software. Relevant structures in electron micrographs were pseudo-colored by a transparent overlay and indicated in the figure legends.

### Statistical Analysis

Measurements were taken from distinct samples and not measured repeatedly, with the exception of **Extended Data Fig. 3g,4i**, where mitochondrial kinetics were replotted for comparison with mitolysosome kinetics. The number of biological replicates and statistical tests are detailed in the figure legends for each experiment. No statistical methods were used to predetermine sample sizes, but the chosen sample sizes are similar to those reported in previous publications (for example, Misgeld et al., 2007; Brill et al., 2016). Comparative datasets were acquired and analyzed in a blinded manner. Samples were not randomized during data collection, due to constraints in animal availability and the assignment of mice to experimental groups according to the given genotypes. Statistical analysis was performed using GraphPad PRISM software (Version 8). Data were tested for normal distribution using the D’Agostino-Pearson normality test. In datasets which passed this test, significance was determined by a two-sided student’s *t*-test. Otherwise, significance was tested by using a two-sided Mann-Whitney-U test. A two-tailed Wilcoxon signed rank test was used for paired analysis. A one-way or two-way ANOVA test with multiple comparisons was used to compare multiple (normally distributed) groups with each other; Data that were not normally distributed were analyzed with a Kruskal-Wallis-test with Dunn’s multiple comparisons test. A two-sided χ²-test was used for contingency analyses. p values < 0.05 were considered significant and marked by ‘‘*’’; p values < 0.01 were indicated by ‘‘**’’ and < 0.001 by ‘‘***.’’ Boxplots show median, interquartile range and whiskers 95th percentiles. Due to the wide data range of mitochondrial transport rates, violin plots were used to represent the entire data distribution and median values. Results written in the main text and in barplots indicate mean ± s.e.m.

### Animal husbandry

Transgenic mouse lines were interbred with C57BL/6J mice on regular intervals (The Jackson Laboratory; Stock #000664). Mice were kept in individually ventilated cages (IVC, Techniplast, Euro-Standard III, at 20 °C, group-housed at max. 5 adult mice per cage) at a 12-h-light-dark-cycle with dry food (ssniff), water *ad libitum* and enrichment. Mice were kept in a maximum-barrier facility with quarterly Felasa monitoring (status stable for more than 10 years). All animal experiments were approved by the Regierung von Oberbayern (Sachgebiet 54) and were in accordance with the national regulations.

### Data Availability

The datasets generated during and/or analysed during the current study are available from the corresponding author on reasonable request. *Thy1*-mito-paGFP and *Thy1*-mito-Dendra mouse lines are available from the Corresponding Author upon request.

## EXTENDED DATA TABLES

**Extended Data Table 1.**
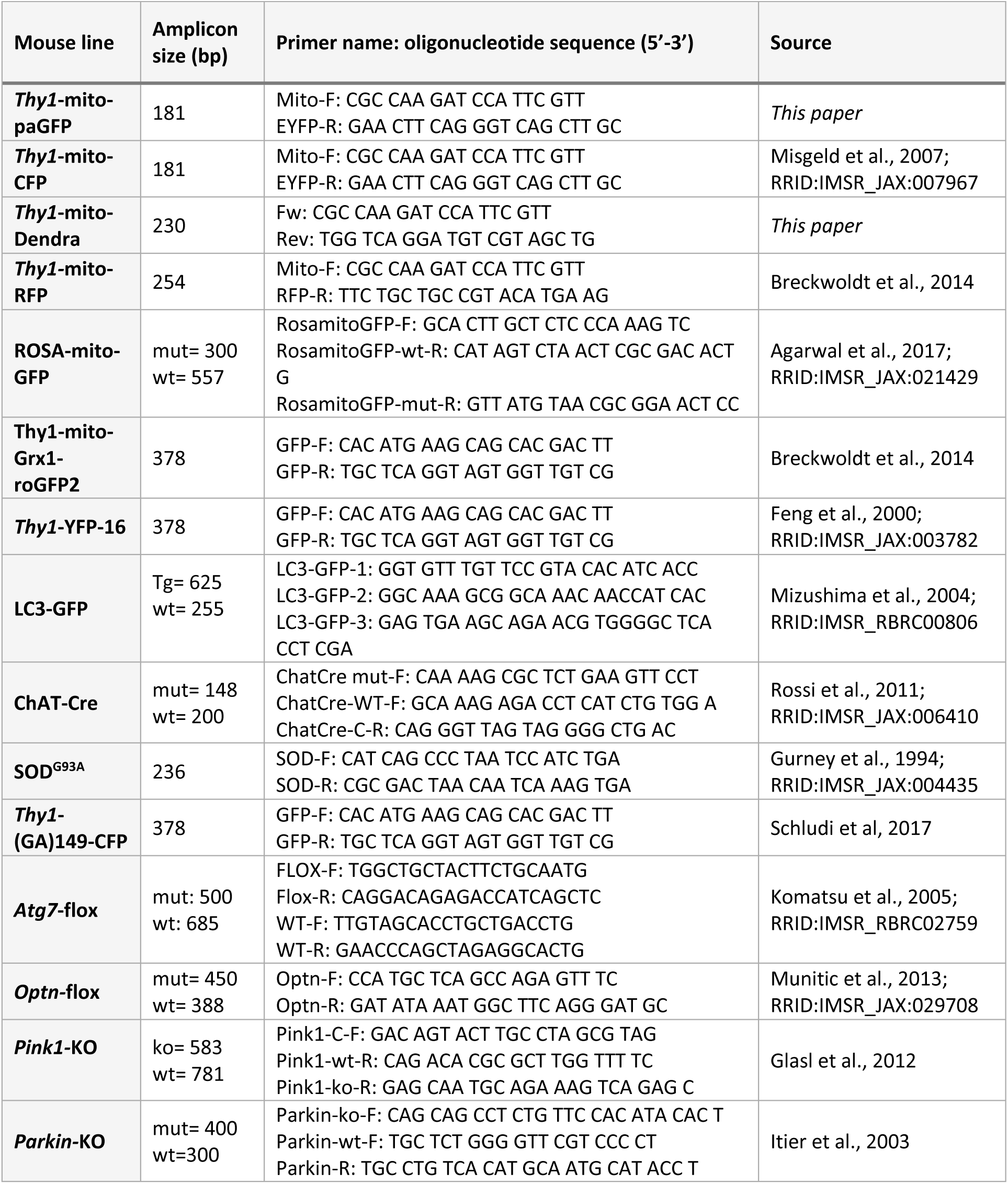
Mouse lines and genotyping primers.

**Extended Data Table 2.** Summary of expression patterns of the *Thy1*-mitopaGFP and *Thy1*-mitoDendra mouse lines.

| Line | Spinal cord |  | Retina |  |  | Cortex | Cerebellum |  |  | Hippocampus |  |  |  | Expression Level |
| --- | --- | --- | --- | --- | --- | --- | --- | --- | --- | --- | --- | --- | --- | --- |
|  | MN | DRG | RG | AC | BC |  | GC | MF | PC | DG | CA1 | CA2 | CA3 |  |
| *Thy1-mitopaGFP;N877* | + | + | + | subset | – | + | – | + | + | + | + | + | + | bright |
| *Thy1-mitoDendra;4* | + | + | + | subset | – | subset | – | + | + | – | – | + | – | bright |

